# Deubiquitinase USP1 influences the dedifferentiation of mouse pancreatic β-cells

**DOI:** 10.1101/2022.10.14.512247

**Authors:** Meenal Francis, Preethi Sheshadri, Jyothi Prasanna, Anujith Kumar

## Abstract

Diabetes is a metabolic disease caused majorly due to loss of insulin secreting β-cells. Along with apoptosis, recent reports revealed dedifferentiation to be the added reason for the reduced β-cell mass. The Ubiquitin Proteasome system comprising of E3 ligase and deubiquitinases (DUBs) control several key aspects of pancreatic β-cell functions. The role of deubiquitinases in orchestrating the dedifferentiation process in several cancers have been well deciphered, but its role in dedifferentiation of pancreatic β-cells remains elusive. In this study, screening for key DUBs that regulate dedifferentiation, identified USP1 to be specifically involved in the process. Inhibition of USP1 either by genetic intervention or small molecule inhibitor ML323 restored epithelial phenotype of β-cells, but not with inhibition of other DUBs. Conversely overexpression of USP1 was sufficient to dedifferentiate β-cells, even in absence of dedifferentiation inducing cues. Mechanistic insight showed USP1 to probably mediate its effect via modulating the expression of Inhibitor of Differentiation (ID) 2. Further, in an *in vivo* streptozotocin (STZ) induced dedifferentiation mouse model system, treatment with ML323 rescued the hyperglycaemic state. Overall, this study assigns a novel role to USP1 in dedifferentiation of β-cells and its inhibition may have a therapeutic application of reducing the β-cell loss during diabetes.

## INTRODUCTION

Diabetes is a metabolic disease presented with a condition of hyperglycaemia owing to the weaning of pancreatic β-cells. Autoimmune destruction of β-cells and β-cell stress due to systemic resistance to insulin are the major reasons for the loss of pancreatic β-cells (Mellitus 2005, Eizirik et al., 2020). Recent theories divulge in accounting to loss of β-cells to multiple cellular attires such as transdifferentiation and dedifferentiation (Moin et al., 2019, Efrat 2019). The latest “Anna Karenina Model” suggests dedifferentiation to be a transitory fate change depending on the stage of the disease (Nimkulrat et al., 2021). Several cell-based lucrative avenues to restore β-cells have been postulated. Transplantation of cadaver donor islets or β-cells derived from differentiation of pluripotent stem cells are ideal cell based strategies, however, they are fraught with shortage of donors and inefficient differentiation respectively (Shapiro et al., 2000, Sutherland et al., 2004). These hurdles could be overcome by formulating a protocol that enhances *in vitro* differentiation or find an approach to enhance endogenous β-cell mass either by blocking transdifferentiation or dedifferentiation.

The *in vitro* culture of islet β-cells is an extremely difficult task owing to the non-proliferative disposition of islet cells and unaccountable fibroblasts like cells emerging from the islet clusters (Kaiser et al., 1988, Hayek et al., 1989, Hayek et al., 1995). In a later study, serum free cultures of β-cells indicated the ability of the cells to shuttle between functional epithelial and mesenchymal fates (Gershengorn et al., 2004). There have been multiple experimentally proven works including lineage tracing studies showing the dedifferentiation process to be occurring in β-cells and proved to be one of the major reason in the manifestation of diabetes (Russ et al., 2008, Russ et al., 2009, Talchai et al., 2012, Xiao X et al., 2017). Dedifferentiation process involves epithelial β-cells transforming to a proliferative mesenchymal cell-like phenotype with non-functional progenitor state (Nimkulrat et al., 2021). Cellular signatures also transit from loss of mature β-cell specific markers such as MAFA and PDX1 to transcribing forbidden genes such as NGN3 and ALDH1A3 and even pluripotent genes (Bensellam et al., 2018). Molecular detailing showed transcription factor forkhead box protein O1 (FOXO1) to translocate to the nucleus under metabolic stress conditions and inhibit the matured β-cells undergoing reprograming to the progenitor state. FOXO1 also rescues key parameters related to diabetes like decreased β-cell mass and function (Talchai et al., 2012, Kitamura et al., 2005, Fiori et al., 2013, Jazurek-Ciesiolka et al., 2019). Injury models such as pancreatic ductal ligation or pancreatectomy conditions have been shown to drive the dedifferentiation of β-cells and the mechanism has been attributed to the signalling of TGFβ secreted by the immune cells (Xiao X et al., 2017). Moreover, adult β-cells during this condition begins expressing EMT gene Snail2 (J Michael Rukstalis et al., 2007).

The homeostasis of transcription factors (TFs) regulating dedifferentiation are in-turn regulated either at transcriptional level or by targeting them to Ubiquitin Proteasome System (UPS) or Lysosomal Autophagy Pathway (LAP) (López-Avalos et al., 2006, Petroski 2008, Hartley et al., 2009, Sun-Wang et al., 2021, Banno et al., 2016, Voutsadakis et al., 2012, Tsubakihara et al., 2018). More than 60% of cellular proteins are subjected to proteasome mediated degradation. The UPS involves ligases (E1, E2 and E3) that facilitate various degrees of ubiquitination resulting in mono, oligo or poly ubiquitinated forms of proteins. In general, proteins ubiquitinated at K48 are targeted to proteasomal degradation (Lingbeck et al., 2003, Nandi et al., 2006). In contrary, deubiquitinases (DUB’s) are set of enzymes which deubiquitinate targeted proteins and thereby salvage them from proteasomal degradation (Ventii et al., 2008, Estavoyer et al., 2022). Several events occurring during pancreatic development or dedifferentiation of β-cells are governed by the UPS. The TF PDX1 which is important for the development of pancreatic epithelial cells is targeted to proteasomal degradation by E3 ligase PDX1 C-terminus–interacting factor-1 (Pcif1). Deletion of FBW7 ubiquitin ligase in adult pancreas drives ductal cells towards β-cells. E3 ligase Mind bomb1 (Mib1) targets Notch receptor Delta to proteasomal degradation and regulates the pancreatic proximal-distal axis patterning of pancreas (Claiborn et al., 2010, Sancho et al., 2014, Itoh et al., 2003, Horn et al., 2012).

The role of DUBs in the context of dedifferentiation of cancer cells are well documented. SNAIL1, a key regulator of dedifferentiation is stabilized in esophageal squamous cell carcinoma, breast cancer cells and lung carcinoma cells by set of DUBs including OTUB1, DUB3, and USP37 respectively (Zhou et al., 2018, Wu et al., 2017, Cai et al., 2020). DUB3 has also been shown to deubiquitinate SLUG and TWIST, the other two essential regulators of dedifferentiation (Lin et al., 2017). Recent few reports provided information on direct impact of DUBs on β-cell physiology (Francis et al., 2022). CLEC16A-RNF41 and USP8 forms a tripartite complex, wherein CLEC16A is an E3 ligase that ubiquitinates RNF41 and USP8 is the DUB that deubiquitinates RNF41. The fine tuning of RNF41 protein level dictates the mitochondrial quality in pancreatic β-cells and facilitates insulin secretion (Pearson et al., 2018, Pearson et al., 2018). Accumulation of toxic misfolded IAPP (Islet amyloid polypeptide) oligomers is one of the reason behind β-cell dysfunction and this is attributed majorly to failure in deubiquitination of IAPP by UCHL1 which further results in aggregation of polyubiquitinated forms (Costes et al., 2011). Similarly, OTUB2 has been shown to deubiquitinate TRAF6 which enhances the survival activity of NFκB in β-cells (Li et al., 2010). MCPIP1 is a peculiar protein which possess both DUB and RNase like activities and is responsible in shielding β-cells from cytokine toxicity (Tyka et al., 2019). Despite these initial insights, comprehensive information about DUBs in dedifferentiation of β-cells largely remains elusive and identifying the key DUBs that regulate dedifferentiation beams to be a resilient proposition to elucidate a novel therapeutic approach.

In the present study, we aimed to investigate and identify key DUBs involved in dedifferentiation of pancreatic β-cells. Screening for DUBs revealed USP1 to be the probable candidate and loss and gain of function further reiterated the importance role of USP1 in dedifferentiation. Administration of small molecule inhibitor of USP1 in a dedifferentiation diabetic mouse model reduced the blood glucose levels and maintained the glucose tolerance test (GTT) and islet architecture near to normalcy. Finally, experiments to examine the downstream effector of USP1 revealed ID2 to be the downstream protein responsible for the dedifferentiation process.

## MATERIALS AND METHODS

### Cell Culture

MIN6 and HEK293T3 cells were maintained in DMEM HG media (Gibco) containing 10% FBS (Thermo Fisher Scientific), 2mM Glutamax (Gibco), Penicillin 100units/ml and Streptomycin 100μg/ml (Gibco). For dedifferentiation of MIN6 cells, the cells were trypsinized with 0.25% trypsin (Sigma Aldrich), counted and plated at the rate of 2 ×10^5^ cells/35mm dish and factors added the following day. The cells were cultured with and without 20ng TGFβ (Sino Biologicals) & 50nM proteasome inhibitor (PI) MG132 (Sigma Aldrich) and maintained for 4 days and later harvested for analysis. Handpicked endogenous islets were plated on 10μg/ml fibronectin (Sigma Aldrich) coated wells and maintained in RPMI media (Gibco) containing 10% FBS, 2mM Glutamax, Penicillin 100units/ml and Streptomycin 100μg/ml (Gibco) with or without 20ng TGFβ for 4 days and later harvested for analysis.

### Plasmid isolation

To isolate the plasmids, the plasmid transformed bacterial broth were pelleted by centrifuging at the rate of 5500rpm for 8 minutes. Following manufacturer’s protocol (Machery Nagel NucleoBond XtraMidi) the plasmids were isolated. Briefly, the bacterial pellet was suspended in 8 ml of RES-EF buffer containing 1mg/mL RNase A. Lysis was performed using LYS-EF buffer at room temperature for 5 minutes followed by neutralization using NEU-EF buffer. Centrifuged at 5000rpm for 15min and the supernatant containing DNA was applied onto the columns. After washes the DNA was eluted using ELU-EF buffer and further precipitated using iso propyl alcohol. The DNA pellet was washed with cold 70% ethanol, dried and the pellet re-suspended in appropriate volume of nuclease free water. DNA concentration was measured using ND1000 Nanodrop and further used for transfections.

### Transfection and transduction

HEK293T3 Cells were plated at the rate of 3 ×10^6^ cells per 60mm dish a day prior to transfection. Transfection agent X-tremeGENE (Merck) was added to optiMEM media (Gibco) at the rate of 1:4 ratio of plasmid concentration, vortexed and incubated for 10min at room temperature. To the transfection mix, the defined concentration of viral packaging plasmids VSV-G and GAG-POL for overexpressing Flag-HA-USP1 (Addgene, #22596) and pLV-TetO-ID2 (Addgene plasmid #70764) and PMD2G and PsPAX2 packaging plasmid for expressing USP1 shRNA and ID2 shRNA, were added and incubated for 30min at room temperature. Following which the mixture was added to the HEK293T3 cells and incubated at 37ºC in CO2 incubator. The culture supernatant containing the viral particles were collected after 48hrs and 72 hrs and were concentrated by incubating in PEG with a final concentration of 50% on a rocker at 4 ºC overnight and centrifuged at 1600rpm for 1hr at 4 ºC. The viral palette was suspended in 200μl media and added to respective cells for transduction. Post 24hrs 1μg/ml of puromycin (HiMedia) was added to respective cultures for selection. Transduced cells were further used for respective experiments. List of plasmids used for the study are provided in Table 3 supplementary material.

### Mouse endogenous islet isolation

All animal experiments were conducted post approval by the Animal Ethics Committee of Manipal Academy of Higher Education (MAHE) (IAEC/KMC/70/2020)). All animals were handled in accordance with the national ethical norms and they were maintained in a facility compliant to national regulations. Pancreas from 2 mice were isolated and digested for 20 min at 37°C in a shaker incubator in digesting enzyme liberase (Sigma Aldrich). The digest was neutralized with 1X HBSS (Sigma Aldrich) and the pellet was subjected to Dextran (Sigma Aldrich) based gradient centrifugation and the top most layer was separated and isolated islet clusters were handpicked using a Nikon SMZ1500 dissection microscope.

### Transcript Analysis

RNA was isolated from cultured cells using Tri®Reagent (Sigma Aldrich) in accordance with manufacturer’s protocol. Briefly, monolayer cultures were homogenized with TRI®Reagent to the mixture. 200μl/ml chloroform was added vortexed for 10 seconds and centrifuged at 12000g for 10min at 4ºC. The aqueous layer was collected and RNA was precipitated by adding 500μl of isopropanol per ml TRI®Reagent and incubated at room temperature for 10min, then centrifuged at 12000g for 10min at 4°C. The RNA pellet obtained was washed with 75% ethanol by centrifuging at 7500g for 5min at 4ºC. After which the supernatant was discarded and the pellet air-dried and suspended in DNAse/RNase free water. RNA was quantified using Naodrop and 1μg RNA was used for cDNA synthesis using commercial cDNA preparatory kit (Takara 6110A). For gene analysis the cDNA was amplified using gene specific primers by qPCR with Takara 2X SYBR Green Mix (Takara RR420A) and analyzed in real time PCR machine (7500 Applied Biosystems). Gene expression was normalized to the expression of housekeeping gene and the relative fold change was calculated using the 2 -ΔΔCt method. List of primers is provided in Table 2 of supplementary material.

### Flow Cytometry

Cells to be analyzed were trypsinized and washed with 1X PBS and fixed by suspending in 4% Paraformaldehyde (PFA) and incubating at 4^0^C overnight. For flow cytometry analysis, the cells were counted and equal number of cells distributed into flow tubes, washed and permeabilized with 1X BD Permwash (BD). The cells were incubated with primary antibody overnight, followed by washes and incubation with secondary antibody (Primary and Secondary antibody dilution provided in Table 1 Supplementary material) for 1 hr again washed and re-suspended in FACS buffer for analysis using BD LSR II or BD FACS Caliber.

### Immunofluorescence

Cells were fixed with 4% PFA and incubated at 4°C. For the experiment, the cells were washed with 1X PBS, then permeabilized with 0.2% Triton X-100 in PBS for 15 minutes at RT, blocked in blocking buffer containing 3% BSA in 1X PBS Solution at RT for 1hr. Antibodies were diluted in blocking buffer and cells were incubated with the primary antibody at 4°C overnight. The cells were subsequently washed with 0.05% PBST and stained with secondary antibody (Primary and Secondary antibody dilution provided in Table 1 Supplementary material) for 1 hr at RT and counterstained with DAPI (1:10000) (Life Technologies) for 1 minute and observed under fluorescent microscope Nikon Eclipse TE2000U.

### SDS PAGE and Western Blotting

Cells were harvested and lysed with ice-cold RIPA lysis buffer with freshly added protease inhibitor cocktail (Sigma Aldrich, Bangalore). The lysate was spun down at 12000 rpm for 15min and the supernatant collected and stored in −20ºC until further use. 100μg of protein was denatured by heating at 95 ºC with protein loading buffer and resolved using 10% SDS-Polyacrylamide gel. The resolved proteins were transferred onto charged PVDF membrane (Millipore) using semi-dry blotting apparatus. The membrane was blocked using 3% skimmed milk in 1x TBST. The blots were probed with primary antibody and incubated overnight at 4ºC on a rocker. Following day, the blots were washed trice with 1X TBST and incubated with Horse Radish-Peroxidase (HRP)-conjugated secondary antibody (Primary and Secondary antibody dilution provided in Table 1 Supplementary material) for 1hr at room temperature on a rocker. The blots were then washed trice with 1X TBST. The blots were developed using Western Bright HRP substrate (Advansta) and imaged on LI-COR C digit blot scanner.

### Dithizone (DTZ) staining

10mg/ml DTZ (Sigma Aldrich) solution was added onto freshly handpicked mouse endogenous islets and incubated for 3-5 min. The stain removed and islets washed with 1X HBSS and phase contrast coloured pictures of red stained islets were captured on Nikon Eclipse TE2000U.

### Hematoxylin and Eosin staining (H & E)

The fixed tissues were embedded in paraffin and 5μm sections were stained by H & E staining. The sections were deparaffinised with Xylene washes, rehydrated with gradient alcohol washes, the slides were stained with hematoxylin for 1 minute and post a 70% alcohol wash stained with Eosin stain for 1 minute. The slides were then processed through graded alcohols and xylene and mounted with DPX and subsequently observed using a phase contrast microscope Nikon Eclipse 80i.

### Immunohistochemistry

Paraffin embedded sections were deparaffinized by placing the slides in a 65ºC hot plate for 30seconds to 1 minute followed by xylene washes and rehydrated with gradient alcohols. The sections were subjected to antigen retrieval by 20min incubation in boiling in 10mM citrate buffer pH 6.0. Blocking with 5% normal serum + 1% BSA in 0.05M Tris Hcl buffer pH 7.5 for 1hr. Washed in Tris Hcl buffer and added primary antibody and incubated overnight. Following day washed in Tris Hcl buffer twice and incubated with fluorescent secondary antibody (Primary and secondary antibody dilution provided in Table 1 Supplementary material) overnight and counter stained with DAPI, mounted with mounting media and observed under Nikon TE80i.

### Blood Glucose tests

Mice was restrained in a tube with tail left out. With a lancet needle prick a droplet of blood was drawn and applied onto glucometer strips and blood glucose concentration was obtained through the glucometer reading (Accu-Chek) in mg/dl.

### Glucose tolerance test (GTT)

16 hours starved mouse were injected intra-peritoneally with glucose (HiMedia) 2g/Kg body mass and blood glucose levels monitored at 0 min, 15^th^ min, 30^th^ min, 60^th^ min and 120^th^ min. The readings were graphically represented in line graph.

### Insulin positive area

c-peptide positive area (cm^2^) in 15 random histological sections of pancreas isolated from animals of respective condition were totalled and represented graphically as fold change compared to control.

### STZ mouse diabetic model and ML323 treatment

With single hand restrain mice starved for 6 hrs were administered intra-peritoneally with 50mg/kg body weight of streptozotocin (Sigma Aldrich) for 6 consecutive days. Along with normal and STZ treated conditions, two conditions of ML323 (Sigma Aldrich) treatment were maintained, a) daily treatment of ML323 6 days post STZ treatment and b) treatment of ML323 beginning 3-days prior to STZ treatment.

### Statistical Analysis

Student’s t-test was used to analyse the difference between the test cells and its respective controls. The values with p<0.05 were considered statistically significant indicated as * and those with p<0.001 were considered highly significant indicate as **.

## RESULTS

### USP1 is upregulated during dedifferentiation of pancreatic β-cells

To identify the key DUB involved in dedifferentiation of β-cells, we generated *in vitro* dedifferentiation model system by subjecting MIN6 cells to TGFβ and Proteasomal Inhibitor (PI) treatment (Takkunen et al., 2006, Dhawan et al., 2016). The cells showed reduced epithelial marker *E-Cadherin* and a corresponding decrease in *insulin* and a moderate increase in mesenchymal marker Snail1 at the transcript level (Supplementary Figure 1a). Flow cytometry and Western blot confirmed the observation by showing reduced levels of E-CADHERIN and PDX1 levels (Supplementary Figure 1b and 1c), cumulatively exhibiting a dedifferentiated state. In this model system, the expression level of multiple DUBs was analysed and identified an increased expression of USP1 compared to other DUBs (Figure 1a and b). To confirm the observation, the transcript analysis was performed along with other DUB *USP30* and observed an enhanced expression of *USP1*, but not the USP30, in TGFβ +PI treated condition compared to untreated cells (Figure 1b). The upregulation of USP1 was consistent at the protein level as observed by flow-cytometry analysis (Figure 1c). To validate the elevated expression of USP1 in a dedifferentiated condition, we isolated primary islets from the mice and authenticated the identity of insulin producing cells by DTZ staining, immunofluorescence of c-peptide (Supplementary Figure 2a, b and c) and transcript analysis of *Insulin, Nkx6.1* and *E-cadherin* (Supplementary Figure 2d). The primary islets cultured on fibronectin coated dish displayed the emanating fibroblast like cells from the islets (Supplementary Figure 3a) and transcript analysis showed down regulation of pancreatic β-cell specific markers *Insulin, Nkx6.1, Pdx1* and epithelial marker *E-Cadherin* and increased expression of mesenchymal markers *Snail1, Snail2* and *Twist* compared to the isolated islets (Supplementary Figure 3b and c). Subjecting the primary islets to TGFβ treatment showed upregulated expression of USP1, but not the other DUBs, similar to that observed in MIN6 cells (Figure 1 d & e). These results probably signify a strong correlation between USP1 and dedifferentiation process in β-cells.

**Figure 1:**
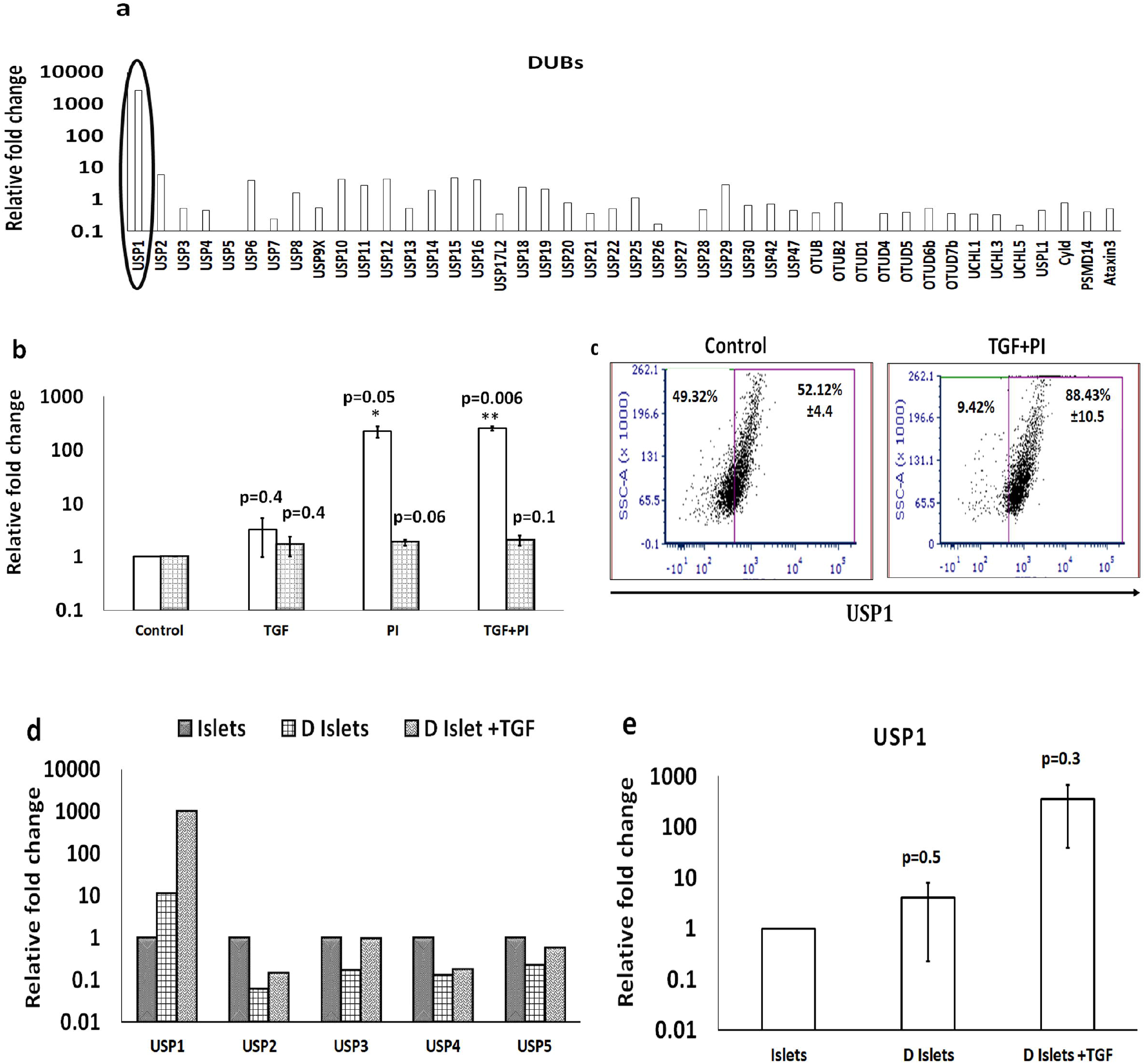
Enhanced expression of USP1 upon mesenchymal transition of pancreatic β cells. (a) MIN6 cells treated with TGFβ+PI were screened for expression of DUBs. Transcript analysis for the expression of various DUBs in MIN6 based model of dedifferentiation compared to untreated control highlighting the expression of USP1. (b) Transcript analysis of USP1 and USP30 in MIN6 cells treated with TGFβ, PI and TGFβ+PI. (c) Dot plots representing flow cytometry analysis for expression of USP1 in MIN6 cells treated with TGFβ +PI compared to control. (d) Transcript analysis for the expression of representative DUBs in *in vitro* dedifferentiated endogenous mouse islets. (e) Transcript analysis of expression of USP1 in dedifferentiated endogenous mouse islets treated with TGF compared to untreated control and whole islets. Data represented as mean ±S.E.M. of 3 sets of experiments *p <0.05 and **p < 0.01.

### USP1 expression is essential for the dedifferentiation of MIN6 cell line

In order to explore the role of USP1 in the dedifferentiation process, we knocked down the expression using USP1 specific shRNA. The USP1 shRNA efficiently inhibited the expression of USP1 which was monitored at the transcript level (Supplementary Figure 4a and b) as well at the protein level (Supplementary Figure 4c). Treatment of MIN6 cells with TGFβ+PI and transduced with scrambled shRNA showed 70% reduction in the expression of *E-Cadherin* and 76% reduction in *insulin* expression at transcript level as assayed by qPCR and E-CADHERIN expression at protein level as analysed by the flow cytometry (Figure 2a and b). Interestingly, transduction of these cells with USP1 specific shRNA showed a complete rescue in the expression of *E-Cadherin* and *insulin* at transcript level and E-CADHERIN at protein level (Figure 2a and b). In corroboration, compared to TGFβ+PI treated cells, there was 30% reduction in expression of mesenchymal gene VIMENTIN in cells transduced with USP1 shRNA (Supplementary Figure 5). However, similar inhibition of USP30 using USP30 shRNA lacked the ability to rescue the expression of E-CADHERIN, denoting the specificity of USP1 in the dedifferentiation process (Supplementary Figure 6). In parallel, similar rescue effect was observed in *in vitro* dedifferentiated mouse primary islets as represented by transcript analysis for the expression of *E-Cadherin* and *Insulin* under conditions of USP1shRNA transduced cells (Supplementary Figure 4d). Previous studies showed USP1 activity could be allosterically inhibited using small molecule ML323 (Qin Liang 2014). Inhibition of USP1 using ML323 in TGFβ+PI treated dedifferentiated MIN6 cells displayed a rescue in the expression of epithelial gene *E-Cadherin* and *Insulin* at transcript level (Figure 2c) and increased expression of E-CADHERIN and decreased expression of VIMENTIN at protein level (Figure 2d and Supplementary Figure 7b). Similar treatment of ML323 in TGFβ treated *in vitro* dedifferentiated mouse primary islets showed rescue in the expression of E-cadherin and insulin (Supplementary Figure 7a). These results indicate the essential role of USP1 in the dedifferentiation of pancreatic β-cells.

**Figure 2:**
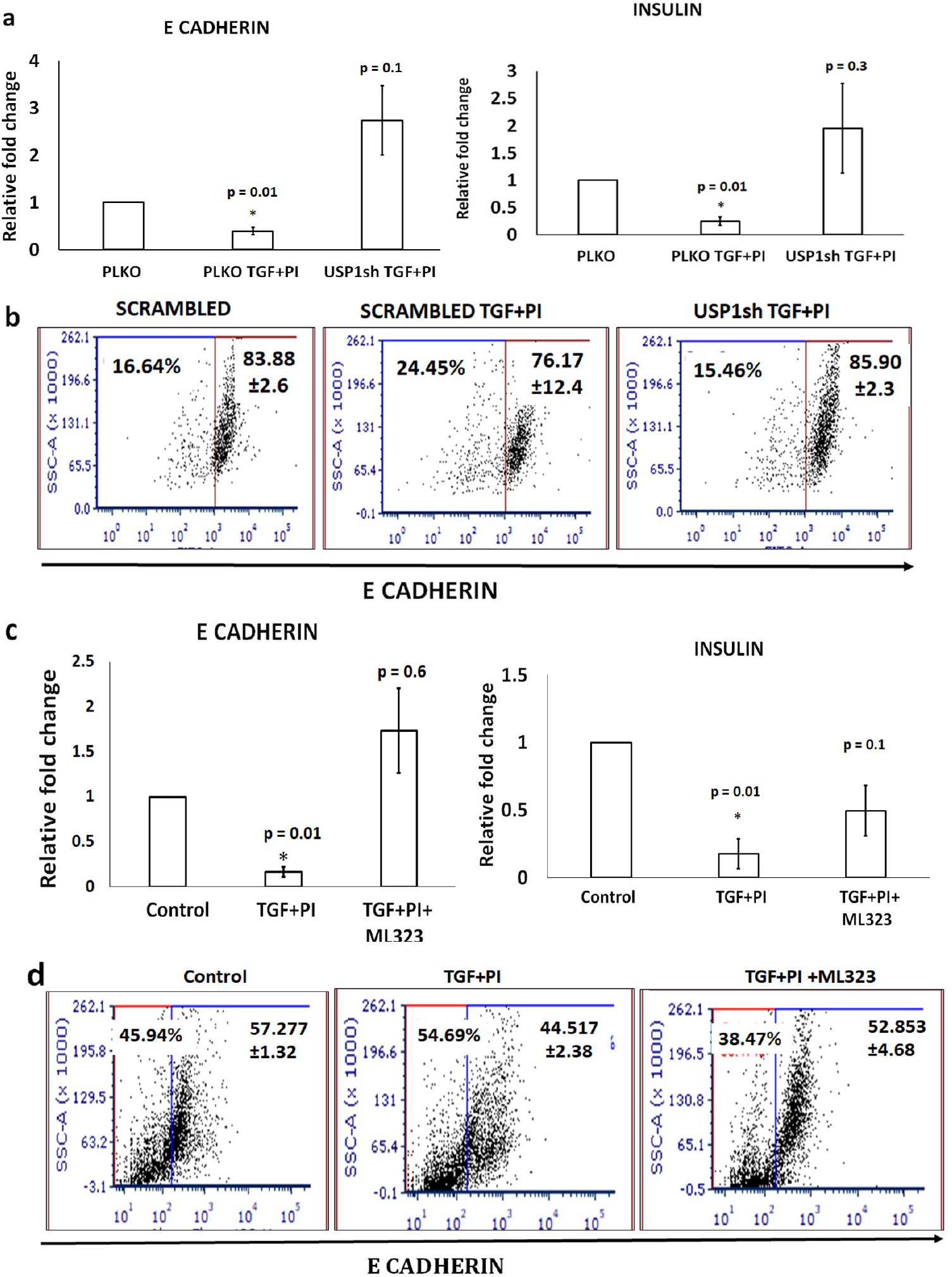
Inhibiting USP1 reverses TGFβ+PI mediated mesenchymal transition. (a) MIN6 cells transduced with USP1sh RNA were treated with TGFβ+PI and analyzed at the transcript level for Insulin and *E-Cadherin*. (b) Flow cytometry analysis for the expression of E-Cadherin in MIN6 cells treated with TGFβ+PI in presence of USP1 shRNA compared to scrambled controls. (c) Transcript analysis of *E-Cadherin* and Insulin in cells treated with TGFβ +PI+ML323. (d) Dot plots representing flow cytometry analysis for the expression of E-Cadherin in cells treated with TGFβ +PI+ML323 or TGFβ+PI compared to untreated control. Data represented as mean ±S.E.M. of 3 sets of experiments *p <0.05 and **p < 0.01.

### Overexpression (OE) of USP1 rescues TGFβ+PI induced dedifferentiation of MIN6 cells

To further confirm the role of USP1 in dedifferentiation of pancreatic β-cells, we analysed the dedifferentiation process in a USP1 overexpressed state. For this, MIN6 cells were transduced with a doxycycline inducible USP1 overexpression construct which upon addition of doxycycline showed enhanced expression of USP1 (Supplementary Figure 8a). Overexpression of USP1 was sufficient to down regulate the expression of *E-Cadherin* and insulin with concomitant increase in the expression of mesenchymal gene *Snail1* compared to TGF+PI treated and untreated controls (Supplementary Figure 8b). To reiterate the specificity of the USP1’s contribution in the dedifferentiation process, USP1 was overexpressed in USP1 shRNA background and analysed for the expression of epithelial genes. Compared to untreated control, TGF+PI treated cells showed 2-fold decrease in the expression of *E-Cadherin* at transcript level and 13 % decrease at protein level. Transduction of USP1 shRNA in this condition rescued the expression of *E-Cadherin* and showed 6-fold increase in expression at transcript level and 12% increased expression at protein level compared to TGF+PI treated cells. Overexpression of USP1 in USP1 shRNA background reversed the expression of *E-cadherin* both at transcript and protein level (Figure 3 a, b, c). To strengthen the observation, instead of shRNA we performed similar experiment in presence of ML323 and observed rescue in the expression of *E-Cadherin* at transcript level and protein levels in cells treated with TGF+PI and ML323 in comparison to cells treated with TGF+PI alone. Overexpression of USP1 in ML323 treated cells reversed the expression of *E-cadherin* and maintained the expression similar to that of TGF+PI treated condition (Figure 3 d, e, f). These results indicate that the increased dedifferentiation observed in β-cells is the consequence of specific upregulation in the expression of USP1 protein.

**Figure 3:**
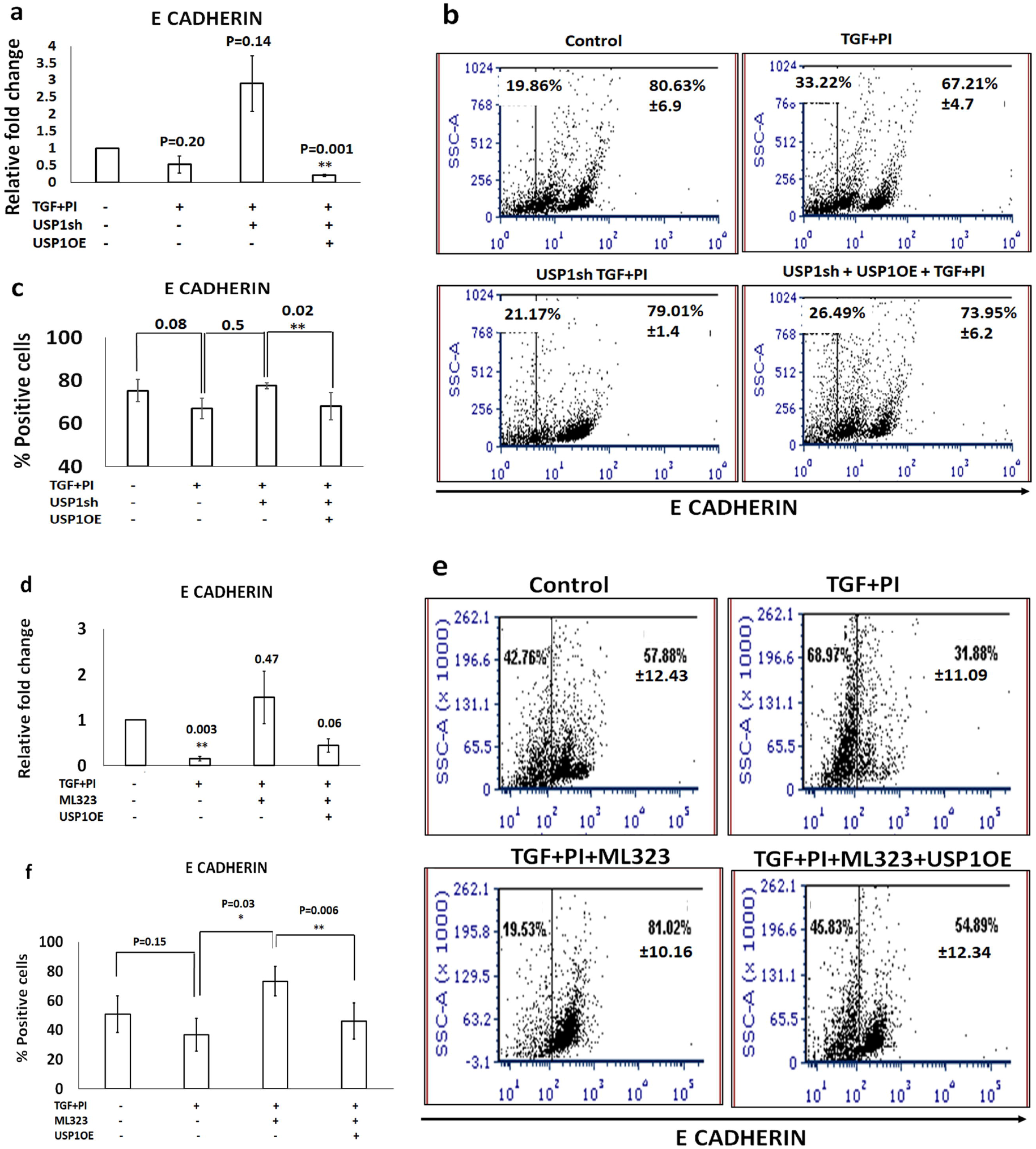
Overexpression of USP1 drives the dedifferentiation of MIN6 cells. (a)Transcript analysis of *E-Cadherin* in USP1sh and USP1 overexpressed cells treated with TGFβ +PI compared to untreated control. (b) Flow cytometry analysis of E-Cadherin in USP1sh and USP1 overexpressed cells treated with TGFβ+PI and compared to untreated control. (c) Quantitative representation of flow cytometry analysis of E-Cadherin expression shown in panel b. (d) Transcript analysis for the expression of *E-Cadherin* in ML323 and USP1 overexpressed cells treated with TGFβ +PI compared to untreated control. (e) Flow cytometry analysis of expression of E-Cadherin in ML323 and USP1 overexpressed cells treated with TGFβ+PI compared to untreated control. (f) Quantitative representation of flow cytometry analysis of E-Cadherin expression shown in panel e. Data represented as mean ±S.E.M. of 3 sets of experiments *p <0.05 and **p < 0.01.

### *In vivo* inhibition of USP1 reversed hyperglycaemia in STZ induced diabetic mouse model

Extending the study to an *in vivo* β-cell dedifferentiation model, we followed the protocol designed by Sachs et. al., 2020, wherein β-cell dedifferentiation was established by administering mice with intra peritoneal injections of STZ for 6 consecutive days. In the mice experiments, the following experimental set up was formulated: a) control mice N=3, b) STZ mice N=3, c) ML323 treatment prior to STZ injection N=3 mice and d) ML323 treatment after STZ injection N=3 mice. Transcript analysis of pancreas in STZ mice showed increased expression of USP1 and decreased expression of glucose transporter isoform GLUT2. The mice treated with USP1 inhibitor prior to STZ treatment showed decreased expression of USP1 and rescued the expression of GLUT2 (Figure 4 a and b). Blood glucose levels were monitored on alternate day points up to day 35. Blood glucose of mice treated with STZ showed an average of 400-500mg/dl. Interestingly, the mice treated with ML323 either 3 days before or 3 days after STZ injection showed reduced blood glucose level with an average of 200-250mg/dl (Figure 4 c). An intraperitoneal GTT was performed after 30 days in mice that received ML323 and observed better clearance of glucose compared to STZ treated and ML323 untreated mice (Figures 4d). Histology of the pancreas of diabetic mice treated with ML323, untreated diabetic and normal animals were analysed by Haematoxylin and Eosin (H&E) staining and immunohistochemical staining for c-peptide. H&E (Figure 4 e, f, g and h) and immunohistochemical staining studies showed rescue in islet architecture in pancreatic sections of mice treated with ML323 compared to STZ treated mice (Figure 4 i-p). Similar rescue in insulin area was also observed in mice treated with ML323 (Figure 4q). Clustering the entire observation, we propose that diabetic animals treated with ML323 sustain functionality of β-cells in the prevailing diabetic conditions probably by inhibiting dedifferentiation.

**Figure 4:**
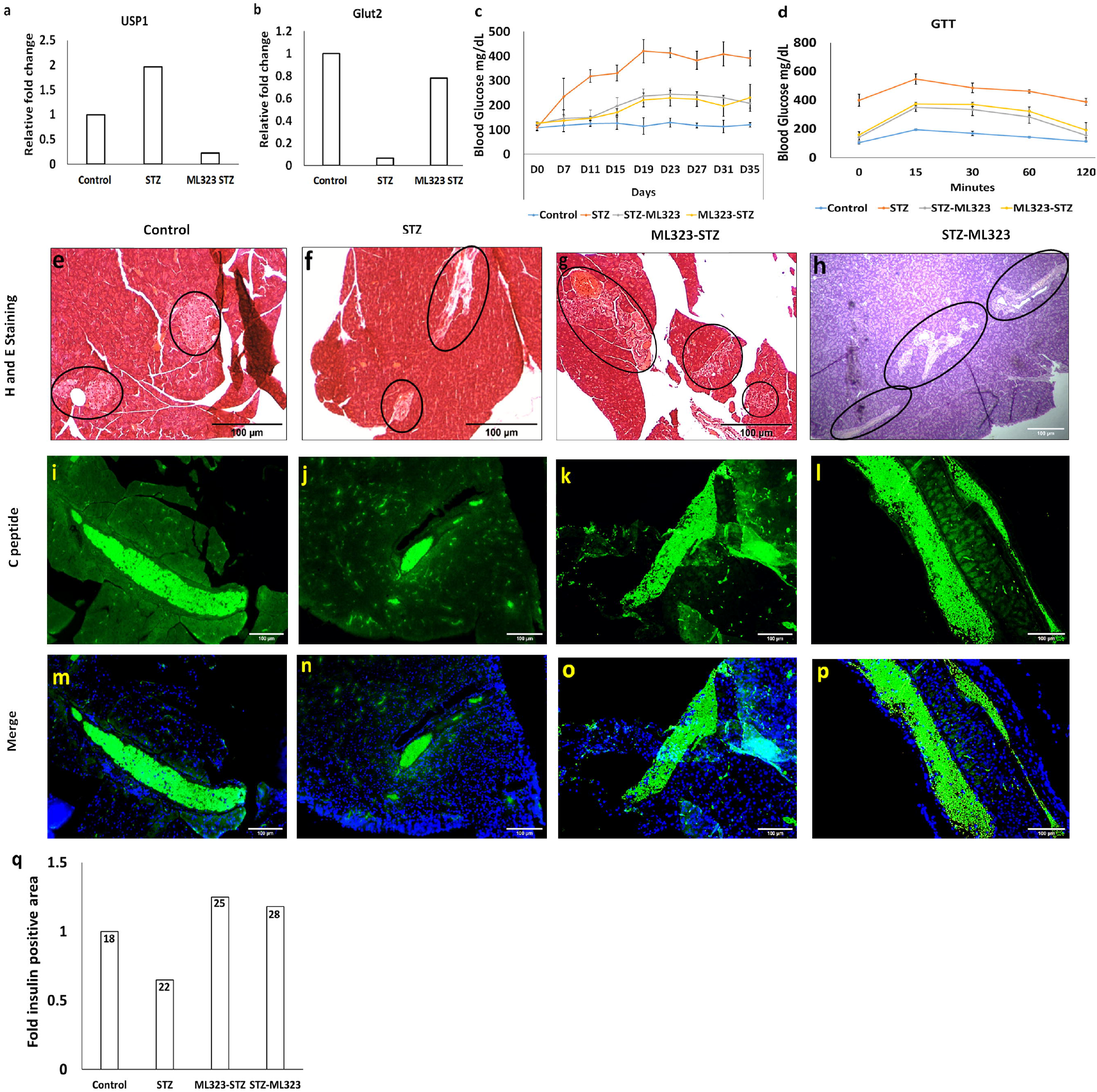
USP1 inhibitor ML323 rescues hyperglycaemia in STZ induced diabetic mice model. (a and b) Transcript analysis for the expression of *USP1* and *Glut2* in the pancreas of STZ and ML323 treated animals compared to untreated controls. (c) Graph representing blood glucose levels across day points of diabetic animals treated with STZ and ML323 compared to untreated animals. (d) Blood glucose levels measured upon glucose tolerance test on day 30 of diabetic animals treated with STZ and ML323 compared to untreated animals. (e,f,g,h) Haematoxylin and eosin stained sections of pancreatic tissue derived from STZ(f), ML323 injected 3 days prior to STZ treatment (ML323-STZ) (g) and ML323 injected 3 days after STZ treatment (h) animals compared to untreated animals (e). (i,j.k.l) Immunohistochemical analysis of c-peptide in pancreatic sections obtained from diabetic animals treated with ML323 (j and k) and untreated diabetic (l) and normal control (i) with respective merge images (m,n,o and p). Data represented as mean ±S.E.M. of 3 sets of experiments. (q) Graph indicating c-peptide positive area of each condition represented as fold change compared to control.

### ID2 is the downstream effector of USP1 mediated dedifferentiation of β-cells

In the prospects of understanding the underlying mechanism of USP1 mediated dedifferentiation of β-cells, we sought after the available information in the literature and found ID family proteins to be the substrates of USP1 in certain metastatic conditions (Williams et al., 2011, Wrighton et al., 2011). This intrigued us to investigate whether ID2 participate in the USP1 mediated dedifferentiation process in the β-cells. Analysis at the transcript level for the expression of ID family genes, ID1, ID2 and ID3 in dedifferentiated islet cells treated with TGF-β compared to untreated controls revealed the upregulation of ID2 compared to ID1 and ID3 (Figure 5a). Further, we looked into the expression of ID family genes in a USP1 inhibited condition in the presence of USP1shRNA in primary islets. Transcript analysis for the expression of ID1, ID2 and ID3 in dedifferentiated islets transduced with USP1sh RNA and treated with TGF-β showed ID2 to be highly down modulated compared to other ID members (Figure 5b). This indicated ID2 protein to be responding to the modulation in USP1 expression level.

**Figure 5:**
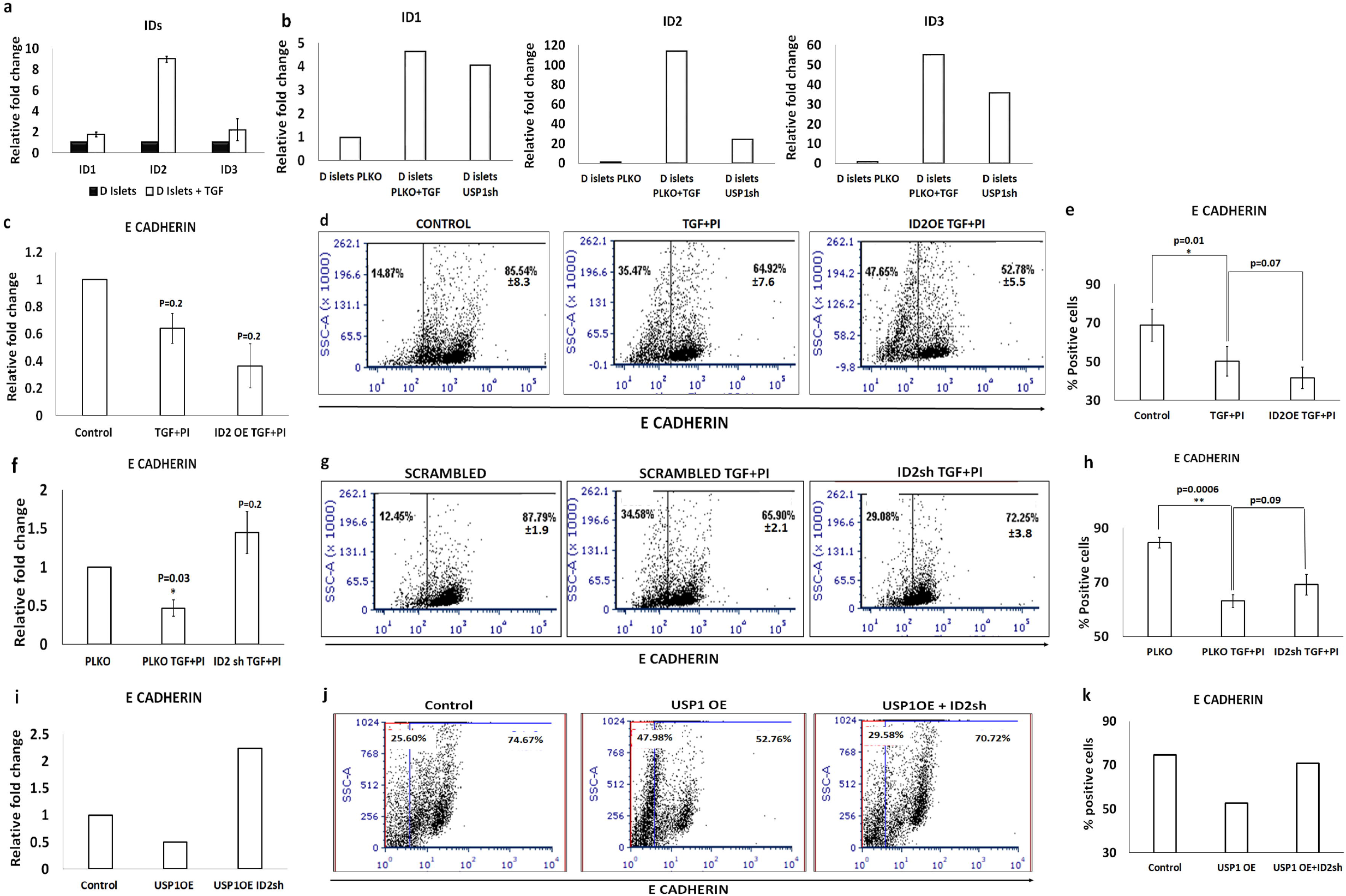
Inhibition of ID2 rescues the USP1 mediated dedifferentiation of β cells: (a) Transcript analysis of ID family genes in TGFβ treated islets compared to untreated counterpart. (b) Transcript analysis of the expression of *ID1, ID2* and *ID3* in the presence of USP1sh RNA in TGFβ treated islets compared to scrambled control. (c) Transcript analysis of the expression *E-Cadherin* in MIN6 cells transfected with ID2 overexpressing construct and treated with TGFβ+PI compared to vector transfected and untreated control. (d) Flow cytometry analysis for the expression E-Cadherin in MIN6 cells harbouring ID2 overexpression construct treated with TGFβ+PI compared TGFβ+PI treated alone and untreated controls. (e) Graphical representation of the flow cytometry analysis indicating percentage positive cells of panel d. Transcript (f) and flow cytometry (g) analysis for the expression of E-Cadherin in MIN6 cells transduced with ID2shRNA and treated with TGFβ+PI compared to scrambled TGFβ+PI treated and scrambled untreated controls. (h) Graphical representation for the flow cytometry analysis indicating percentage positive cells of panel g. Transcript (i) and flow cytometry (j) analysis for the expression of E-Cadherin in MIN6 cells transduced with ID2shRNA in a USP1 overexpression background treated with TGFβ+PI compared to USP1 OE alone and vehicle control. (k) Graphical representation for the flow cytometry analysis indicating percentage positive cells of panel j. Data represented as mean ±S.E.M. of 3 sets of experiments *p <0.05 and **p < 0.01.

To further reveal whether ID2 be the downstream effector of USP1 in regulating the dedifferentiation process, we strategized to perform ID2 loss and gain of function in presence or absence of USP1 in MIN6 cells. Doxycycline induced overexpression of ID2 in cells cultured in presence of TGF+PI reduced the expression of *E-Cadherin* by 2-fold at transcript level and by 12% at protein level compared to doxycycline untreated cells (Figure 5 c, d and e). Inhibition of ID2 expression by ID2 specific shRNA rescued the TGF+PI mediated inhibition of E-Cadherin expression. There was ~3.2-fold increase in the expression of *E-Cadherin* at transcript level and ~8% increase at protein level compared to TGF+PI treated scrambled control (Figure 5 f, g and h). To correlate the relation between USP1 and ID2, we knocked down ID2 in presence of USP1 overexpression condition and analysed for the expression of *E-Cadherin* in MIN6 cells. In cells overexpressing USP1, ~ 2-fold decrease in the expression of *E-Cadherin* at a transcript level and 22% at protein level compared to vector control was observed. Inhibition of ID2 in this USP1 overexpression condition by transducing ID2 specific shRNA reversed the expression of *E-Cadherin* by ~4.5 fold at transcript level and 18% at protein level compared to USP1 overexpression and scrambled shRNA transduced condition (Figure 5 i, j and k). In conclusion, these experiments reveal the activation of USP1 during dedifferentiation process coincide with ID2 expression and indicate the likeliness of ID2 being a downstream effector of USP1 during β-cells dedifferentiation.

## Discussion

Dedifferentiation is a cardinal process playing a dogmatic role during pancreatic β-cell development as well in metabolic diseases like diabetes (Cole et al., 2009, Gouzi et al., 2011, Talchai et al., 2012). Several experimental evidences showed the occurrence of β-cell dedifferentiation which could be one of the major reason for depleted mass in hyperglycaemic state (Jesus et al., 2021, Ibrahim et al., 2021). Retaining *in vivo* β-cell mass by avoiding dedifferentiation is an ideal alternative strategy to combat β-cell loss. In the present study we investigated the role of DUB USP1 and observed its role in *in vitro* and *in vivo* dedifferentiation of pancreatic β-cells. Inhibition of USP1 facilitated in retaining the epithelial phenotype and thereby properties of β-cell. Attempts to understand the mechanism shed light on ID2 to be the probable downstream substrate of USP1 during this process. The outcome of the present study helps in better understanding the process of dedifferentiation in β-cells and small molecule mediated inhibition of USP1 takes a step closer to sustain the β-cell mass.

Several studies showed various signalling pathways to participate in the dedifferentiation process. Recent study by Xiao et al., showed the TGFβ secreted by M2 macrophages to be responsible for the dedifferentiation process in of β-cells in a ductal ligated mouse model system (Xiao X et al., 2017). Due to lesser number of DUBs being encoded by mouse genome compared to E3 ligases, screening for DUBs in dedifferentiation process proved more feasible. Considering the evidence of TGFβ in dedifferentiation, we dedifferentiated MIN6 cells line by treating with TGFβ and PI and screened for key DUBs based on their expression level. We detected the transcript levels of *USP1* to be highly up-regulated compared to other DUBs. Further, enhanced expression of *USP1* were also observed in primary β-cells treated with TGFβ. Several DUBs have been identified to enhance the downstream TGFβ activity by stabilizing the intermediates of TGFβ pathway. Interestingly, the deubiquitinase enzyme FAM/USP9X stabilizes Smurf1, the downstream of TGFβ cascade and facilitates in enhancing the TGFβ activity (Xie Y et al., 2013). However, in our screening process, we failed to detect the enhanced expression of any other DUBs other than *USP1.* Inhibition of USP1 either by shRNA or by small molecule ML323 showed similar effect on dedifferentiation process. Previous studies had shown ML323 to inhibit the enzymatic activity of USP1 allosterically (Liang et al., 2014, Dexheimer et al. 2014). However, in our present study we consistently observed the downregulation of USP1 expression at transcript and protein levels also when the cells were treated with ML323. This decrease in USP1 transcript could be probably a secondary effect caused due to the induction of dedifferentiation process rather than the direct effect of ML323 on the transcription of USP1 expression. There are a few recent reports, which showed a direct correlation between USP1’s activity and manifestation of β-cell failure. Inhibition of USP1 was shown to protect β-cells from glucose toxicity induced apoptosis and the mechanistic insights showed USP1 inhibition to enhance the DNA damage response and in turn to facilitate the β-cell survival (Gorrepati et al., 2018). In our study, though we failed to find difference in β-cell survival upon overexpression or inhibition of USP1, our experimental results indicate inhibition of dedifferentiation in USP1 knockdown condition to be the other alternative mechanism involved in retaining β-cell function.

To date, there has been an ongoing debate on whether β-cells per se undergo dedifferentiation or the mesenchymal cells exist from the pre-existing mesenchymal cells, or is it the transdifferentiation of other islet cells (Francis et al., 2022). Though the ambiguity persists till date, several lineage tracing studies including the elegant study by Talchai et. al., showed the β-cell specific knockout of FOXO1 to result in dedifferentiation of β-cells that results in diabetes (Talchai et al., 2012). Similarly, Mak et al., demonstrated β-cell dedifferentiation to be part of EMT and indicated miR-7 to be the key player in enhancing the expression of mesenchymal genes leading to EMT (Mak et al., 2019). All this collective information portrayed dedifferentiation process to exist in β-cells which triggers loss of β-cell mass and imply to be one of the major causatives of diabetes. However, in our present study, though we observed the occurrence of dedifferentiation in *in vitro* cultured primary islet cells, we hesitate to rule out the contribution of other cells of islets as we considered our analysis to the entire pool of dedifferentiated cells emanating from the cultured islets and not from the fractionated homogenous β-cells. Since larger part of islets is populated with β-cells (60-70%), we still strongly consider the role of USP1 in the process of dedifferentiation of β-cells. To conclusively ascertain the role of USP1 in dedifferentiation, we adapted an *in vivo* diabetic dedifferentiation model system. Sachs et al., demonstrated streptozotocin-induced diabetes model could be utilized to study β-cell dedifferentiation in mice (Sachs et al., 2020). In the present study, in STZ treated pancreas there was increased expression of *USP1* and reduced expression of matured β-cell gene *Glut2.* Surprisingly, administration of USP1 inhibitor ML323 in STZ-induced dedifferentiation model system showed rescue of blood glucose levels and a rescue in the expression of *Glut2* similar to untreated control. The islet architecture as well the insulin positive area of diabetic mice treated with ML323 showed similar phenotype to that of non-diabetic controls but the untreated diabetic animals showed regressed islet clusters and reduced insulin area. These results convincingly portrayed the significant role of USP1 in *in vitro* and *in vivo* dedifferentiation of β-cells.

Mechanistically, dedifferentiation is controlled by several transcription factors. For instance, hypoxia-inducible factor-1α (HIF-1α) has been shown to facilitate dedifferentiation of β-cells (Liu et al., 2020). Conditional knockout of FOXO1 TF resulted in dedifferentiation of β-cells and has been shown to be the major cause of β-cell failure in Type II diabetes (Talchai C 2012). During unfavourable dedifferentiation inducing conditions, FOXO1 translocates to the nucleus and protects the β-cells by facilitating the expression of matured markers NEUROD1 and MAFA (Kitamura et al., 2005). The other important TFs family that regulates dedifferentiation process is ID family members. Elevated ID proteins are observed in several dedifferentiated primary human malignancies including pancreatic carcinoma and neuroblastomas (Maruyama et al., 1999, Chakrabarti et al., 2013). There are various DUBs which regulate the homeostasis of several proteins that dictate the degree of dedifferentiation process. For instance, homeostasis of NGN3, the TF that ear marks islet precursor cells, has been shown to be dependent on the presence of NOTCH signalling (Wang et al., 2010). DUB USP9x stabilizes the components of NOTCH and thereby increases NGN3 activity which in turn probably increases the dedifferentiation process (Premarathne et al., 2017). An independent study also showed the expression of SNAIL in NGN3 expressing precursor cells which inhibited the maturation to β-cells and in parallel facilitated the dedifferentiation process (Rukstalis et al., 2007). Recent study in an independent cellular system showed USP1 to deubiquitinate SNAIL1 and enhance its stability (Sonego et al., 2019). In the present study, with the goal of identifying the mechanism by which USP1 executes its role in dedifferentiation, we considered the previously available information from literature which showed the deubiquitination of ID proteins by USP1 to be essential for maintenance of mesenchymal phenotype in osteosarcoma cells (Williams et al., 2011). In the process of testing whether the similar mechanism holds true in dedifferentiation of β-cells, we observed enhanced expression of ID2 compared to other ID family members in dedifferentiated MIN6 cells. Overexpression of ID2 was sufficient to induce the dedifferentiation of MIN6 cells even in absence of USP1, and inhibition of ID2 even in USP1 overexpression condition was adequate to retain the epithelial gene expression. These results indicated ID2 to be the probable downstream candidate of USP1 and deubiquitination of which probably enhances dedifferentiation of β-cells.

Endogenous β-cells undergoing dedifferentiation mediated changes in *in vitro* culture has crippled the long-term expansion of β-cells. Therefore, it is prudent to have a better understanding of the dedifferentiation process in context of β-cells to retain functional cells in *in vitro* as well for *in vivo* therapeutic applications. Considering the outcome of the present study, altering USP1 activity could be envisaged as an alternative means to sustain *in vitro* β-cell culture as well to inhibit the *in vivo* β-cell loss in diseased state.

## Acknowledgements

This study was supported by grants from Indian Council of Medical Research, India (No. 5/4/5-5/Diab/21-NCD-III) to A.K. Flag-HA-USP1 was a gift from Wade Harper (Addgene plasmid# 22596) and pLV-TetO-ID2 was a gift from Hubert Schorle (Addgene plasmid #70764). We thank AK lab members for their suggestions and encouragement.

## Conflict of interest

All the authors declare no conflict of interest. The authors of the above noted article, Anujith Kumar and Meenal Francis have filed certain aspects of this work for patenting.

## Notes

### Competing Interest Statement

The authors have declared no competing interest.

