## Supplemental material for "Deubiquitinase USP1 influences the dedifferentiation of mouse pancreatic β-cells"

### Supplementary data

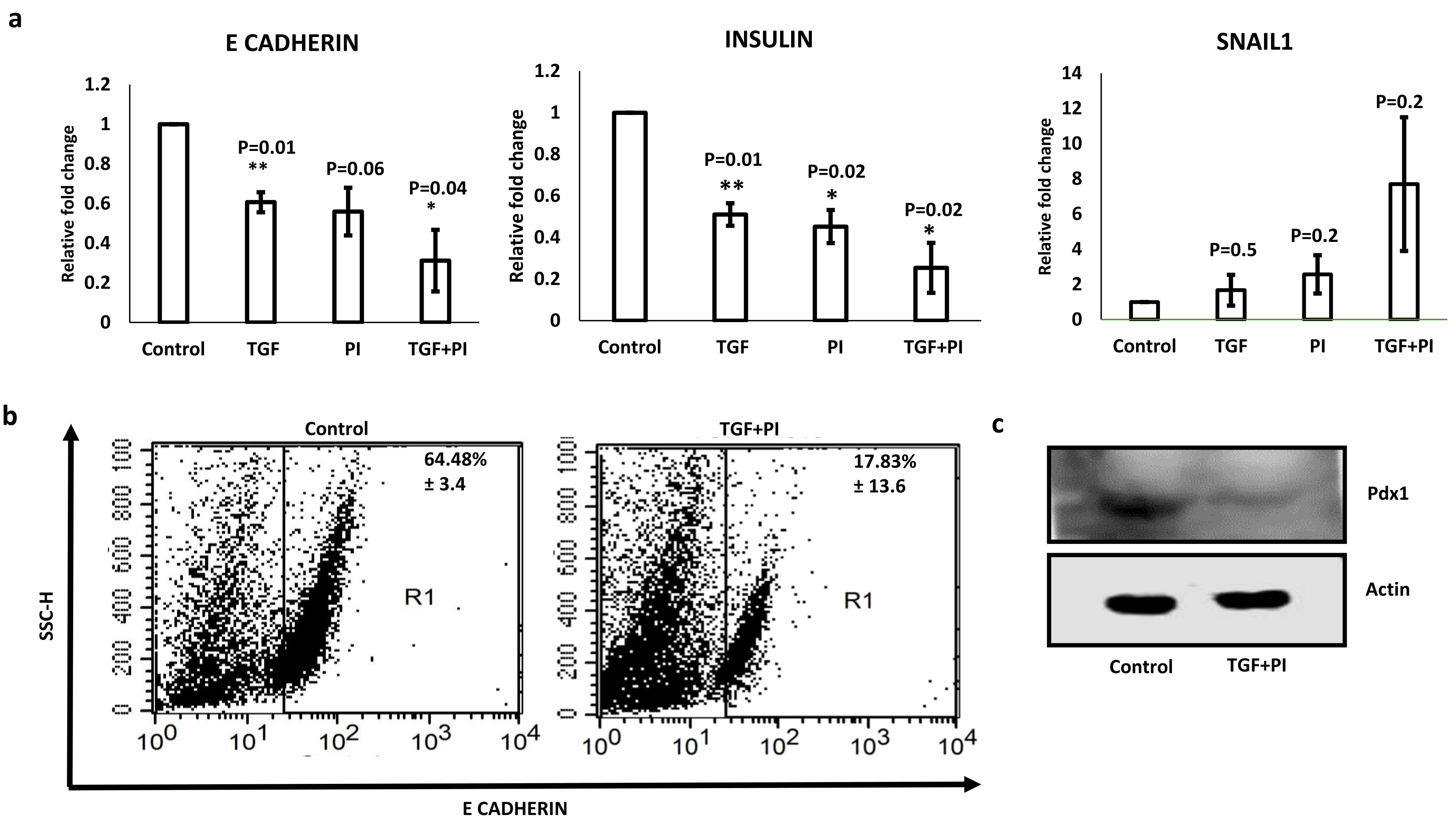

a
E CADHERIN
Relative fold change
1.2
1
0.8
0.6
0.4
0.2
0
P=0.01
**
P=0.06
P=0.04
*
Control
TGF
PI
TGF+PI
INSULIN
Relative fold change
1.2
1
0.8
0.6
0.4
0.2
0
P=0.01
**
P=0.02
*
P=0.02
*
Control
TGF
PI
TGF+PI
SNAIL1
Relative fold change
14
12
10
8
6
4
2
0
P=0.5
P=0.2
P=0.2
Control
TGF
PI
TGF+PI
b
SSC-H
Control
64.48%
± 3.4
R1
TGF+PI
17.83%
± 13.6
R1
E CADHERIN
c
Pdx1
Actin
Control
TGF+PI

**Supplementary Figure 1: Directed dedifferentiation of MIN6 cells towards mesenchymal phenotype upon treatment with TGF and PI**. (a) Transcript analysis for the expression of E-Cadherin, Insulin and Snail1 in MIN6 cells treated with TGF, PI and TGF+PI. (b) Flow cytometry analysis for the expression of E-Cadherin of MIN6 cells treated with TGF +PI compared to untreated control. Data represented as mean ±S.E.M. of 3 sets of independent experiments. *$p<0.05$ and **$p<0.01$ relative to control. (c) Western blots for the expression of Pdx1 compared to house keeping gene actin for MIN6 cells treated with TGF+PI compared to untreated controls.

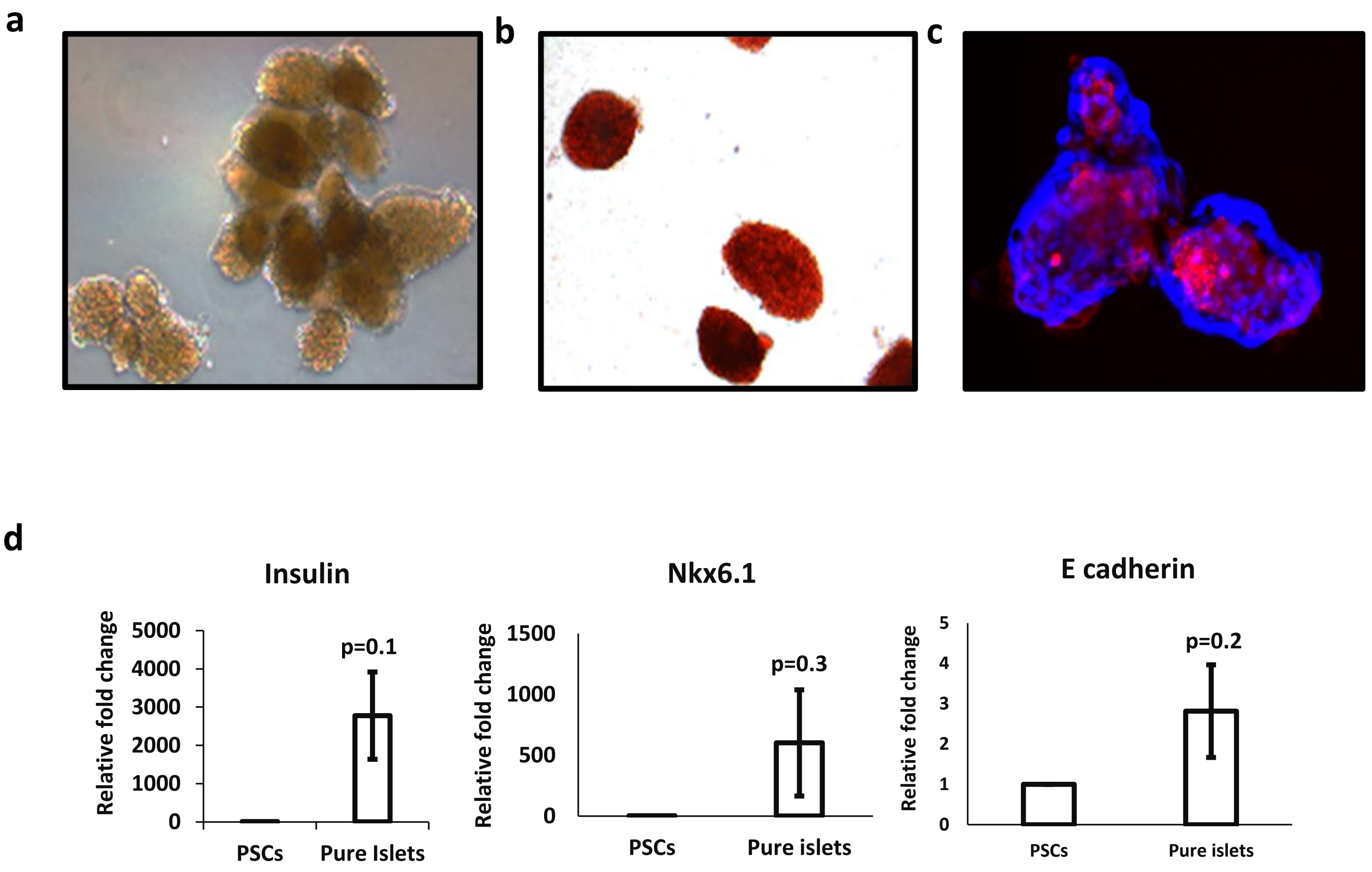

a
b
c
d
Insulin
Relative fold change
5000
4000
3000
2000
1000
0
p=0.1
PSCs
Pure Islets
Nkx6.1
Relative fold change
1500
1000
500
0
p=0.3
PSCs
Pure islets
E cadherin
Relative fold change
5
4
3
2
1
0
p=0.2
PSCs
Pure islets

**Supplementary Figure 2: Characterization of the primary mouse islets fractioned by density gradient.** (a) Phase contrast image of *in vitro* isolated endogenous islets. (b) DTZ stained islets isolated *in vitro*. (c) Immunofluorescence image of primary mouse islets stained for c-peptide (red) and nuclear (DAPI). (d) Transcript analysis for the expression of islet specific markers Nkx6.1 and Insulin and epithelial marker E-Cadherin compared to pluripotent stem cells (PSCs). Data represented as mean ±S.E.M. of 3 sets of experiments $^*p < 0.05$.

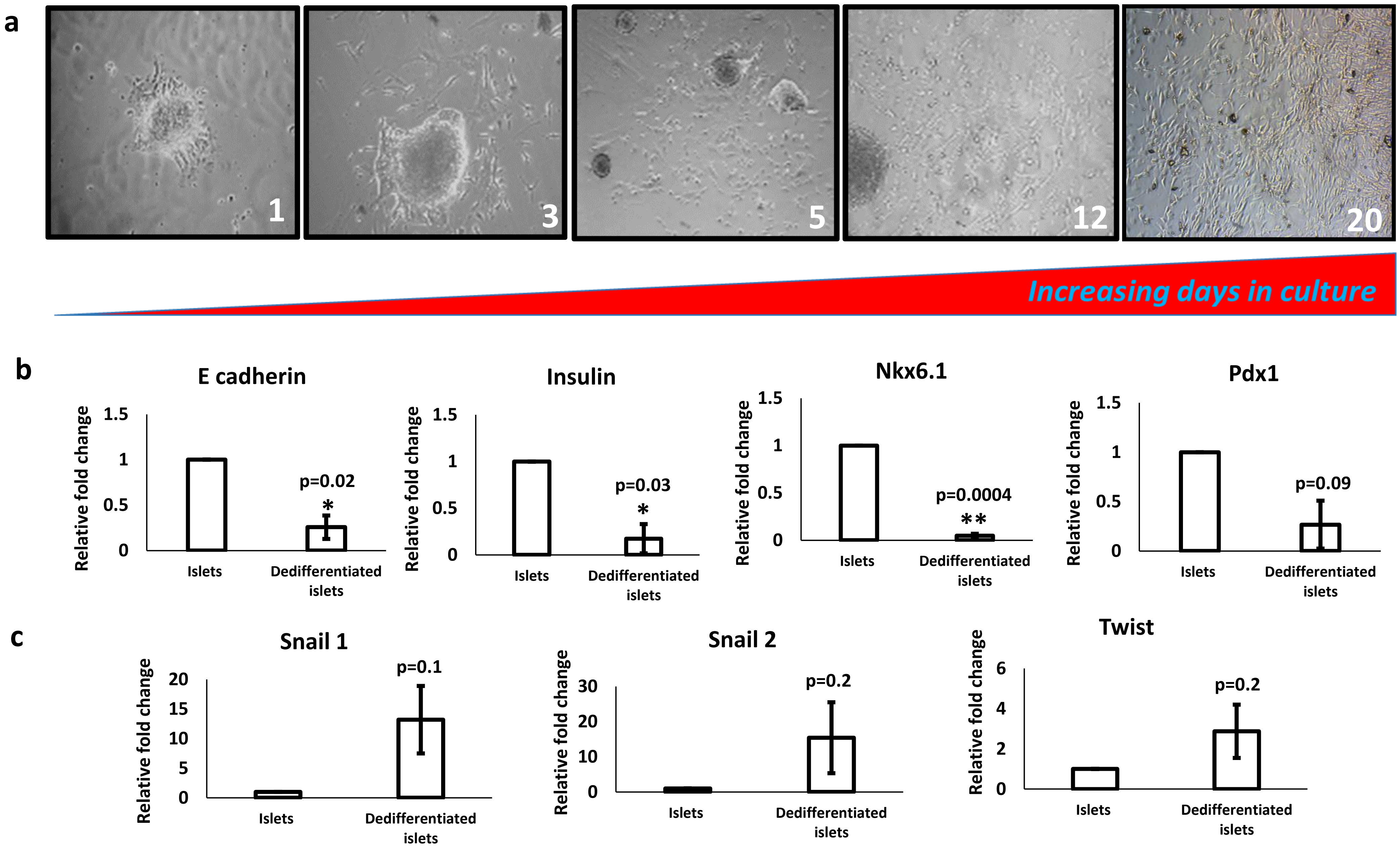
a
1
3
5
12
20
Increasing days in culture
b
E cadherin
Insulin
Nkx6.1
Pdx1
Relative fold change
1.5
1
0.5
0
p=0.02
*
p=0.03
*
p=0.0004
**
p=0.09
Islets
Dedifferentiated islets
c
Snail 1
Snail 2
Twist
20
15
10
5
0
30
6
4
2
p=0.1
p=0.2
p=0.2

**Supplementary Figure 3: Mouse primary islets cultured *In vitro* undergo dedifferentiation.** (a)Phase contrast images of dedifferentiated mouse endogenous islets captured at different day points. (b) Transcript analysis of *in vitro* dedifferentiated islet cells for the expression of pancreatic β-cell specific markers Insulin, Nkx6.1 and Pdx1 and epithelial marker E-Cadherin compared to whole islets. (c) Transcript analysis of *in vitro* dedifferentiated islet cells for the expression of mesenchymal specific markers Snail1, Snail2 and Twist. Data represented as mean ±S.E.M. of 3 sets of experiments. *$p < 0.05$ and **$p < 0.01$.

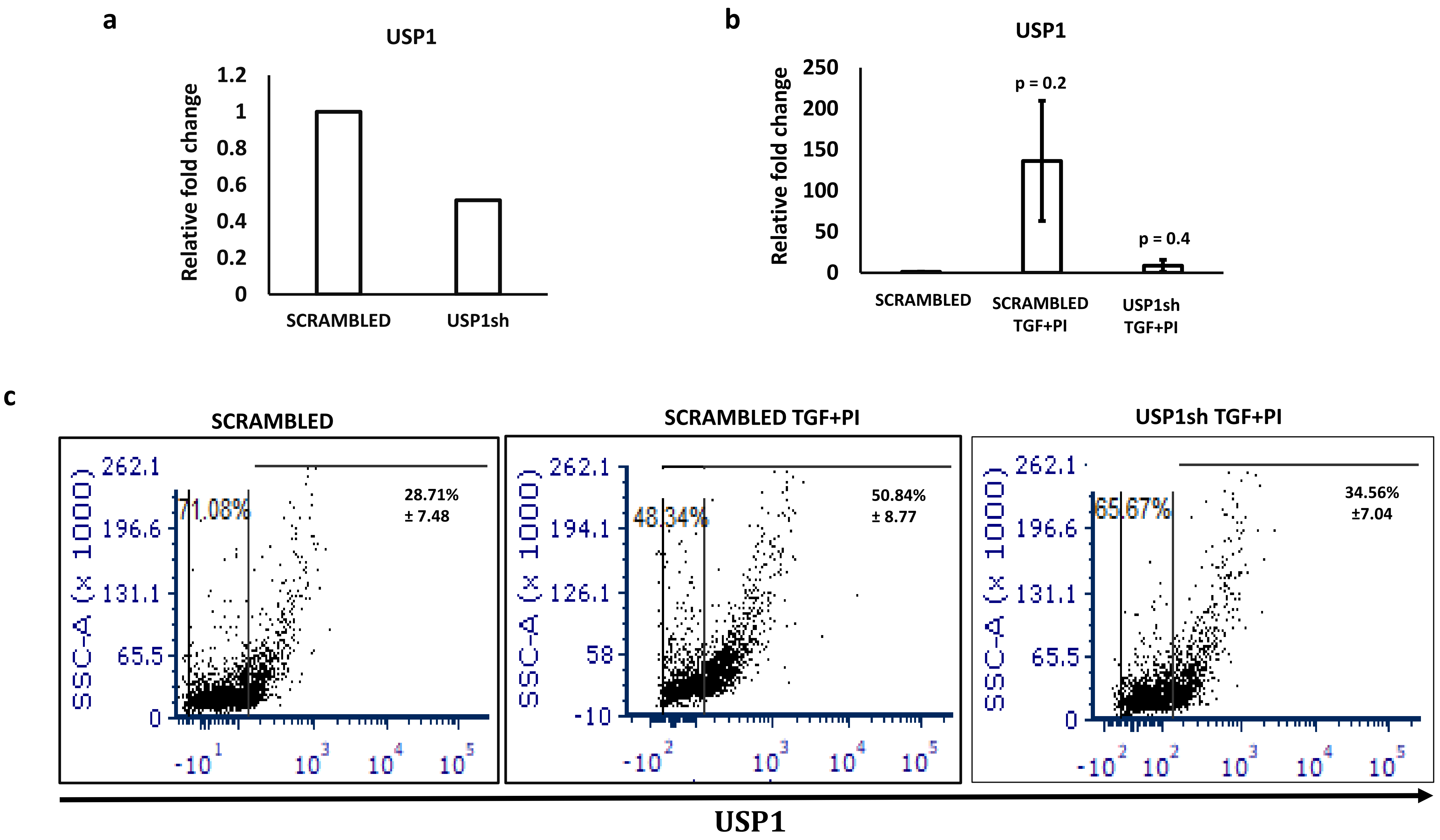
a
USP1
Relative fold change
1.2
1
0.8
0.6
0.4
0.2
0
SCRAMBLED
USP1sh
b
USP1
Relative fold change
250
200
150
100
50
0
p = 0.2
p = 0.4
SCRAMBLED
SCRAMBLED TGF+PI
USP1sh TGF+PI
c
SCRAMBLED
SCRAMBLED TGF+PI
USP1sh TGF+PI
SSC-A (x 1000)
262.1
196.6
131.1
65.5
0
71.08%
28.71% ± 7.48
262.1
194.1
126.1
58
-10
48.34%
50.84% ± 8.77
262.1
196.6
131.1
65.5
0
65.67%
34.56% ±7.04
USP1

d

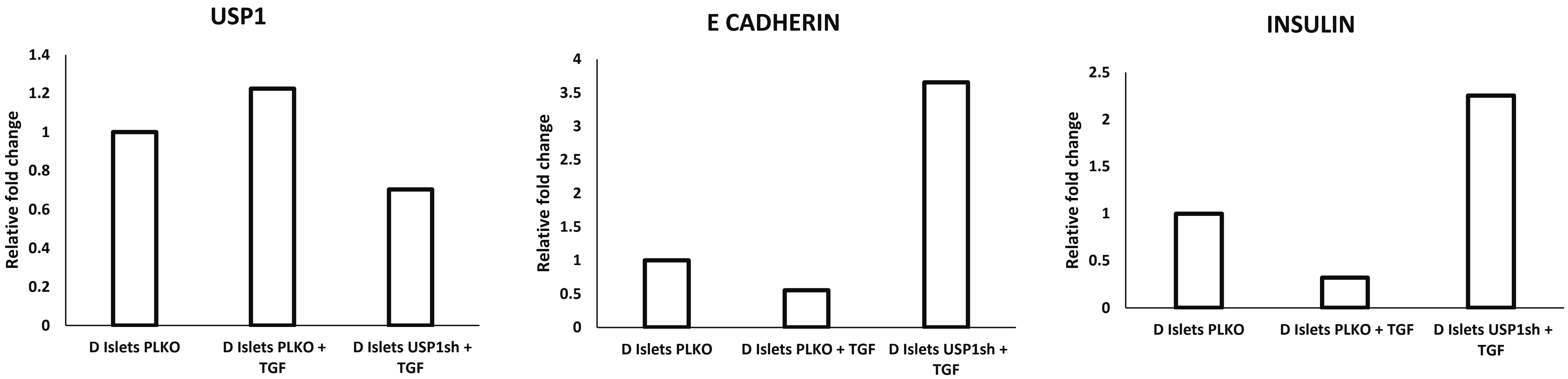
USP1
Relative fold change
1.4
1.2
1
0.8
0.6
0.4
0.2
0
D Islets PLKO
D Islets PLKO + TGF
D Islets USP1sh + TGF
E CADHERIN
Relative fold change
4
3.5
3
2.5
2
1.5
1
0.5
0
D Islets PLKO
D Islets PLKO + TGF
D Islets USP1sh + TGF
INSULIN
Relative fold change
2.5
2
1.5
1
0.5
0
D Islets PLKO
D Islets PLKO + TGF
D Islets USP1sh + TGF

**Supplementary Figure 4: Knockdown of USP1 inhibits the dedifferentiation of islet cells.** (a) Transcript analysis for the expression of USP1 in MIN6 cells transduced with USP1shRNA compared to scrambled controls. (b) Transcript analysis for the expression of  USP1 in MIN6 cells transduced with USP1sh RNA and treated with TGF+PI compared to scrambled control and untreated cells. (c) Flow cytometry analysis for theexpression of USP1 in MIN6 cells treated with TGF+PI and USP1 shRNA compared to scrambled vector control and untreated cells. Data represented as mean ±S.E.M. of 3 sets of experiments. (d) Transcript analysis for the expression of USP1, E-Cadherin and Insulin in dedifferentiated islets transduced with USP1shRNA and treated with TGF compared to scrambled and untreated controls.

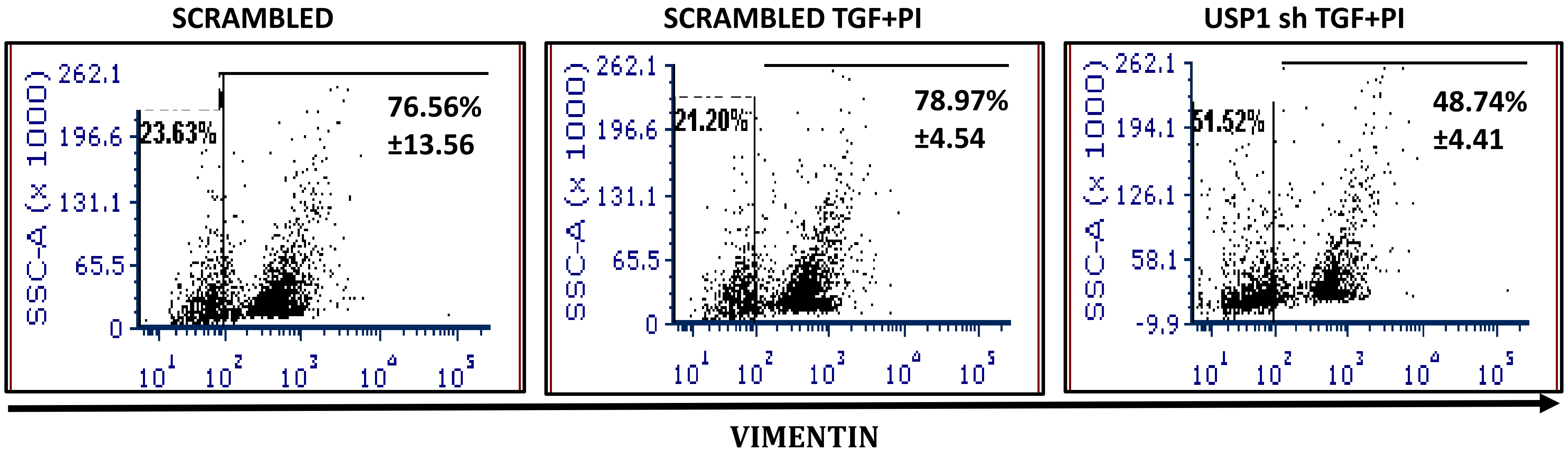


**Supplementary Figure 5: Expression of mesenchymal gene Vimentin correlated with the expression of USP1**. Flow cytometry analysis for the expression of Vimentin in MIN6 cells transduced with USP1shRNA and treated with TGF+PI compared to cells transduced with scrambled shRNA. Data represented as mean ±S.E.M. of 3 sets of experiments.

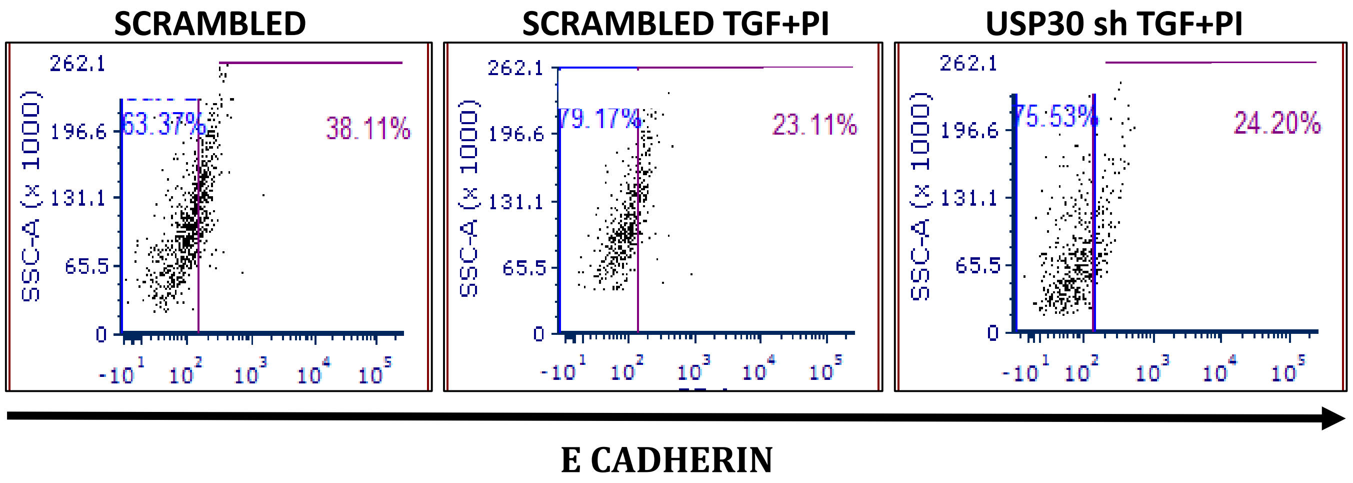


**Supplementary Figure 6: Inhibition of USP30 lacks the ability to rescue epithelial characteristics of MIN6 cells**. Flow cytometry analysis for the expression of E-Cadherin in MIN6 cells transduced with USP30shRNA and treated with TGF+PI compared to cells transduced with scrambled shRNA. Data represented as mean ±S.E.M. of 3 independent sets of experiments.

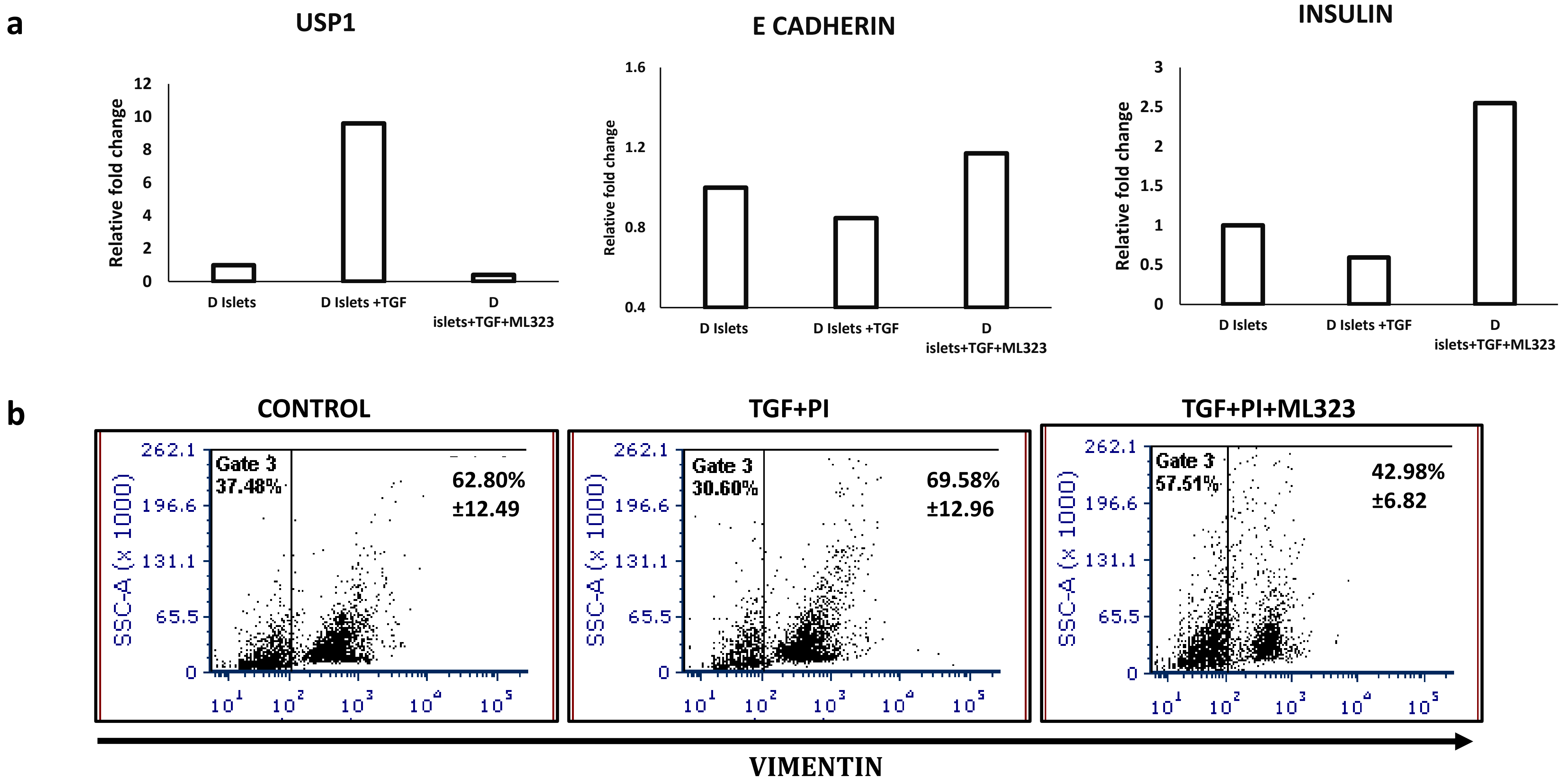


**Supplementary Figure 7: Small molecule inhibitor ML323 inhibits the expression of USP1**. a) Transcript analysis for the expression of USP1, E-Cadherin and Insulin in dedifferentiated islets treated with ML323 and TGF and the expression level compared to scrambled and untreated controls. b) Flow cytometry analysis for the expression of Vimentin in MIN6 cells treated with TGF+PI and cultured in the presence or absence of USP1 inhibitor ML323. Data represented as mean ±S.E.M. of 3 sets of experiments.

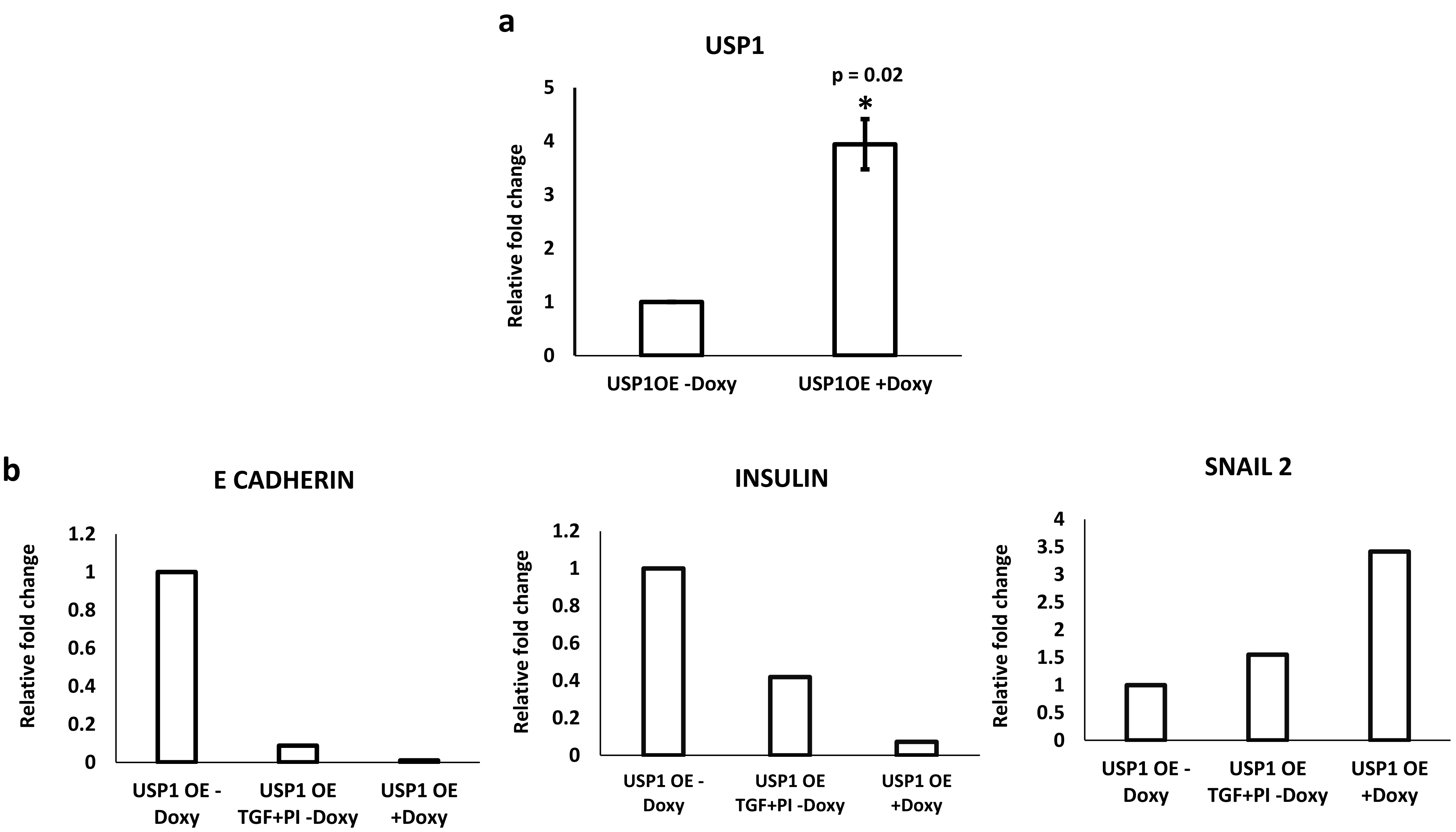
a
USP1
p = 0.02
*
Relative fold change
5
4
3
2
1
0
USP1OE -Doxy
USP1OE +Doxy
b
E CADHERIN
Relative fold change
1.2
1
0.8
0.6
0.4
0.2
0
USP1 OE - Doxy
USP1 OE TGF+PI -Doxy
USP1 OE +Doxy
INSULIN
Relative fold change
1.2
1
0.8
0.6
0.4
0.2
0
USP1 OE - Doxy
USP1 OE TGF+PI -Doxy
USP1 OE +Doxy
SNAIL 2
Relative fold change
4
3.5
3
2.5
2
1.5
1
0.5
0
USP1 OE - Doxy
USP1 OE TGF+PI -Doxy
USP1 OE +Doxy

**Supplementary Figure 8: Forced expression of USP1 is sufficient to dedifferentiate MIN6 cells**. (a) Transcript analysis for the expression of USP1 for MIN6 cells transduced with a doxycycline inducible USP1 overexpression construct compared to un-induced control. Data represented as mean ±S.E.M. of 3 sets of experiments *$p < 0.05$. (b) Transcript analysis for the expression of E Cadherin, Insulin and Snail1 of MIN6 cells transduced with doxycycline inducible USP1 overexpression (OE) construct compared to un-induced cells treated with TGF+PI and untreated controls.

**Supplementary Material: Table 1: List of antibodies used for the study**

| Serial number | Antibody Name | Company | Catalogue Number | Dilution | Technique used |
|---|---|---|---|---|---|
| 1 | Octamer 4 | BD Biosciences, Torreyana Road, San Diego, CA. http://www.bdbiosciences.com/ | BD611203 | 1 : 1000 | Immunofluorosence (IF) |
| 2 | SSEA1 | BD Biosciences, Torreyana Road, San Diego, CA. http://www.bdbiosciences.com/ | BD560079 | 1 : 1000 | IF |
| 3 | Pdx1 | Abcam, http://www.abcam.com/ | ab47267 | IF: 1:200<br>WB: 1: 2000 | IF and Western blots (WB) |
| 4 | C-peptide | Abcam, http://www.abcam.com/ | ab14181 | 1 : 200 | IF |
| 5 | USP1 | Invitrogen | PA5.65023 | 1 : 100 | Flow cytometry (FC) |
| 6 | E cadherin | BD Biosciences, Torreyana Road, San Diego, CA. http://www.bdbiosciences.com/ | 610181 | FC: 1: 100<br>WB: 1:1000 | FC and WB |
| 7 | Actin | Santacruz Santa Cruz Biotechnology, Inc.<br>Dallas, Texas, U.S.A.<br>http://www.scbt.com/ | SC47778 | 1 : 1000 | WB |
| 8 | Sox2 | Millipore, Single Oak Drive, Temecula, CA. http://www.emdmillipore.com/ | MAB4343 | 1 : 1000 | IF |
| 9 | Nanog | Abcam, http://www.abcam.com/ | BD560259 | 1 : 1000 | IF |
| 10 | CXCR4 | Santacruz Santa Cruz Biotechnology, Inc.<br>Dallas, Texas, U.S.A.<br>http://www.scbt.com/ | SC53534 | 1 : 500 | IF |
| 11 | C-peptide | Cell Signalling Technology3 Trask Ln, Danvers, Massachusetts, USA | 45935 | 1 : 500 | IHC |
| 12 | Anti-Rabbit FITC | Invitrogen<br>Thermo Fisher Scientific<br>168 Third Avenue<br>Waltham, MA USA | F2765 | 1 : 500 | FC |
| 13 | Anti-Mouse FITC | Invitrogen<br>Thermo Fisher Scientific<br>168 Third Avenue<br>Waltham, MA USA | A11059 | 1 : 500 | FC |
| 14 | Anti-Mouse HRP | Invitrogen<br>Thermo Fisher Scientific<br>168 Third Avenue<br>Waltham, MA USA | 62-6520 | 1 : 1000 | WB |
| 15 | Anti-Rabbit HRP | Thermo Fisher Scientific<br>168 Third Avenue<br>Waltham, MA USA | 32460 | 1 : 1000 | WB |

| | | | | | |
|---|---|---|---|---|---|
| 16 | Anti-mouse 488 | Thermo Fisher Scientific<br>168 Third Avenue<br>Waltham, MA USA | A21042 | 1 : 1000 | IF |
| 17 | Anti-Rabbit 568 | Invitrogen<br>Thermo Fisher Scientific<br>168 Third Avenue<br>Waltham, MA USA | A10042 | 1 : 1000 | IF |
| 18 | Anti-Mouse 594 | Life Tech<br>Thermo Fisher Scientific<br>168 Third Avenue<br>Waltham, MA USA | A11005 | 1 : 1000 | IF |

**Supplementary Material: Table 2: List of primers used for the study**

| Serial number | Name | Primer sequence | Amplicon size (bp) |
|---|---|---|---|
| 1 | Insulin | F:TGTTGGTGCACTTCCTACCC | 300 |
| | | R:GCTGGTAGAGGGAGCAGATG | |
| 2 | E-Cadherin | F:CGCACTACTGAGTTCCCAAG | 173 |
| | | R:CAAAGCCATGAGGAGAC | |
| 3 | Octomer4 | F:GAGGAGTCCAGGACATGAA | 153 |
| | | R:AGATGGTGGTCTGGCTGAAC | |
| 5 | Snail1 | F:CGGAGTTGACTACCGACCTT | 265 |
| | | R:GGGGTACCAGGAGAGAGTCC | |
| 6 | Snail2 | F:AACATTTCAACGCCTCCAAG | 231 |
| | | R:GCCCCAAGGATGAGGAGTAT | |
| 7 | USP1 | F:CACCAGCGATCCCTCTCC | 148 |
| | | R:TGCCTCTTGAAAGCCCATTA | |
| 8 | USP2 | F:CACACTGTGGGAGAAGAGCA | 112 |
| | | R:GTTTCGAAGACCAGCCAGAC | |
| 9 | USP3 | F:ACTCAGCCAAGTTCCCCAAC | 106 |
| | | R:CCACAGTGGACACTTGAACA | |
| 10 | USP4 | F:GGATGTGGAGACCCAGAAGA | 129 |
| | | R:CAGCTGTCAAAGCCCACATA | |
| 11 | USP5 | F:GCTGTCAGTGTTACCGACGA | 149 |
| | | R:CCACATACTGCTTCCCGAAT | |
| 12 | USP25 | F:GGCAACGACAGGTACATCAG | 105 |
| | | R:CAAACTCAAGGCAATCGCTC | |
| 13 | USP26 | F:CCCAGCCTCCATTTTGTCTC | 109 |
| | | R:GTCCTCACTCTGGTTCACAC | |
| 14 | USP27x | F:CCTTGACCCACACTCCAATAC | 101 |
| | | R:AGCGACGACATCTCACAGAC | |
| 15 | USP28 | F:CCAAAGAGTTAAGGAGCCCAG | 115 |

| | | | |
|---|---|---|---|
| | | R:GTTGTCGTGAGTGAGGTCTATC | |
| 16 | USP29 | F:TGACACCGAAGACATTTGCC | 111 |
| | | R:CAGTGCAAGACAAAAGAGCC | |
| 17 | USP30 | F:ACAGAGAGGAAGAAGCGGAG | 120 |
| | | R:CAGCCACTTGACAAACGCAG | |
| 18 | USP47 | F:ATTTGACCAGGGAAAATGGAG | 148 |
| | | R:AACATCTGAGGCGGTTCAC | |
| 19 | USP7 | F:CACAAGGAAAACGACTGGG | 118 |
| | | R:CATCTGCCTGTACAAAAACTTC | |
| 20 | USP9X | F:ATGGGCTAACGATCTCATTTAC | 120 |
| | | R:TAGCCACACATAGCTCCAC | |
| 21 | USP13 | F:ATCGAAGTATGCCAACAACC | 117 |
| | | R:CCGTCAGTCAGATTCAACC | |
| 22 | OTUB1 | F:CTGATGGCAACTGCTTCTAC | 112 |
| | | R:TCCTCTTTACTCTTGGCAGAC | |
| 23 | USP17L2 | F:GGAAAGCACAGTCTGAGTTG | 100 |
| | | R:CAACTCAGACTGTGCTTTCC | |
| 24 | USP6 | F:GCAACTGGAAATCGAAAGGAC | 144 |
| | | R:TCAAGAAGAAGAGCCCAGAC | |
| 25 | USP8 | F:TCCTGAAGAAGCTATCAGTCC | 108 |
| | | R:ACCACATAGTCCACACTCC | |
| 26 | USP10 | F:AAAATCGTGAGAGATATCCGCC | 123 |
| | | R:TAGATACTCCTCTGCGTCCTCC | |
| 27 | USP11 | F:GCACCTTTCCTGGCTGTATC | 104 |
| | | R:TGGGAGCAGCACATAATCATC | |
| 28 | USP12 | F:TCGGCATTAGAGAAAGAGATTG | 137 |
| | | R:TTTTCCCGAAATGGACGAC | |
| 29 | USP14 | F:CAGAAGAACCCTCTGCTAAAAC | 147 |
| | | R:CAGGCACAGAACGAATACAC | |
| 30 | USP15 | F:CGGGAACATGAACAATGTTGT | 116 |
| | | R:ACAACATTGTTCATGTTCCCG | |
| 31 | USP16 | F:TGAAGCACTACACGACACC | 110 |
| | | R:TGAGCTACAGTACTTGACTTCC | |
| 32 | USP18 | F:TGGAGAAGATGCAGGACAG | 141 |
| | | R:GGTCCTTAGTCAGGTTCCAG | |
| 33 | USP19 | F:ATGCTGCGTTCACAGACAC | 134 |
| | | R:GTACTAGCTGTAGAAGACCACC | |
| 34 | USP20 | F:GAAGTCTTTCTGGAGCAGC | 123 |
| | | R:TTCATCAGCCACAGCAATAG | |
| 35 | USP21 | F:ACTCCTGTGAAGCTGTGAATCC | 133 |
| | | R:ATCTCAAGGTGCAACCGCTC | |
| 36 | OTUD7b | F:CCTTCCAGCTTCCAGATCTCAC | 122 |

| | | | |
|---|---|---|---|
| | | R:AACCCACCAATTCAGACGCC | |
| 37 | UCHL5 | F:GGCTGTATGAACTAGACGGG | 101 |
| | | R:CCTTTTCTCTATTACTGGCCTC | |
| 38 | USPL1 | F:CCGAAGCTTAGATGCACATTAG | 140 |
| | | R:TGGTGTCCACAATGGGAAC | |
| 39 | OTUD6B | F:TATTGTGAACACAGCGGC | 125 |
| | | R:TCGGATACTCTTCACCGAC | |
| 40 | OTUD5 | F:TCAAGCAGATGAAGGAGGAC | 102 |
| | | R:CCATGCAATGCTTTCGAAC | |
| 41 | UCHL1 | F:AGCAGACCATCGGAAACTC | 123 |
| | | R:GCTTCTCCGTTTCAGACAG | |
| 42 | UCHL3 | F:ACAAACCATCAGCAATGCC | 127 |
| | | R:GCTCATTGATACAGACTCCTCC | |
| 43 | USP22 | F:TGTGTCTTCTTTGGCTGTTTC | 102 |
| | | R:GCAGTAAATACCTCCGTACATC | |
| 44 | PSMD14 | F:GTTGATGATTACACCGTCAGAG | 126 |
| | | R:TCCTGTTTGTTTCAGCATATCC | |
| 45 | ATAXIN3 | F:TGGCTCAATTACAGCAAGAAG | 138 |
| | | R:GTGCAAGTTCCTCTCCAATAAG | |
| 46 | USP42 | F:TCCCATGAACACTCCAAGAC | 116 |
| | | R:ACGAACATCGGCTTGATAAC | |
| 47 | OTUB2 | F:GACCAAAGGAGACGGAAAC | 105 |
| | | R:AGCACACGCTCTTTGAAC | |
| 48 | CYLD | F:TTGAAGGTTGGAGAAAGTACAG | 123 |
| | | R:TCCATCCCAGTTGCCAATAG | |
| 49 | OTUD1 | F:TTCAAGGTCTCCTTGCAGTC | 100 |
| | | R:GGTACTTGTTTCTCTGCCTC | |
| 50 | OTUD4 | F:CCCAGTCTCAGAAATCCTCC | 102 |
| | | R:CGTGATCAAAGTCCTCAGCC | |
| 51 | GAPDH | F:AACTTTGGCATTGTGGAAGG | 132 |
| | | R:GGATGCAGGGATGATGTTCT | |
| 52 | Nkx6.1 | F:TCAGGTCAAGGTCTGGTTCC | 211 |
| | | R:CGATTTGTGCTTTTTCAGCA | |
| 53 | Pdx1 | F:ACAGCCCTGAGCTTCTGAAA | 162 |
| | | R:CTGCTGGTCCGTATTGGAAC | |
| 54 | Twist | F:GCTCAGCTACGCCTTCTCC | 297 |
| | | R:CCTCTGGGAATCTCTGTCCA | |
| 55 | Glut2 | F:TTCAGTACAGGACCTGGATTAAGAG | 151 |
| | | R:CTTGAGGTGCATTGATCACAC | |
| 56 | AFP | F:GCCCTACAGACCATGAAACAAG | 149 |
| | | R:GTGAAACAGACTTCCTGGTCCT | |
| 57 | GATA6 | F:CAACACAGTCCCCGTTCTTT | 122 |

| | | | |
|---|---|---|---|
| | | R:TGGTACAGGCGTCAAGAGTG | |
| 58 | Sox17 | F:GATGCGGGATACGCCAGTG | 136 |
| | | R:CCACCACCTCGCCTTTCAC | |
| 59 | Cxcr4 | F:CTTCTGGGCAGTTGATGCCAT | 156 |
| | | R:CTGTTGGTGGCGTGGACAAT | |
| 60 | Foxg1 | F:CTGAGTGTGGACCGGCTG | 213 |
| | | R:CGTGCTGGTCTGCGAAGTC | |
| 61 | Sox1 | F:CTGCTCAAGAAGGACAAGTA | 416 |
| | | R:CTCATGTAGCCCTGAGAGT | |
| 62 | Vegf | F:CTGCTCTCTTGGGTGCACTG | 375 |
| | | R:TTCACATCTGCTGTGCTGTAG | |
| 63 | Brachury | F:TCCCGAGACCCAGTTCATAG | 106 |
| | | R:TTCTTTGGCATCAAGGAAGG | |
| 64 | Rex1 | F:CCAACAGGAAAGTGAATGGTG | 195 |
| | | R:CAAAGTTGGCCATTCTTTAGGA | |
| 65 | Hnf6 | F:CCCTGGAGCAAACTCAAGTC | 256 |
| | | R:GCTGGGAGATGGTGATTTGT | |
| 66 | MafA | F:CGGGACCTGTACAAGGAGAA | 150 |
| | | R:GGCACGTCACAGAAAGAAGTC | |
| 67 | YFP | F:TACCAGTTCCATGGCCAACT | 383 |
| | | R:TCAGCTAATTGAACACCACCA | |
| 68 | ID1 | F:GCAGCATGTAATCGACTACATC | 243 |
| | | R:ACTTTTTTCCTCTTGCCTCC | |
| 69 | ID2 | F:GCATCCCCCAGAACAAGAAG | 228 |
| | | R:CATTCGACATAAGCTCAGAAGG | |
| 70 | ID3 | F:ACTACATCCTCGACCTTCAG | 252 |
| | | R:ATCGAAGCTCATCCATGCC | |
| 71 | USP1 (Human) | GCTCTAAAGGATGAAGCCAATC | 122 |
| | | CTAGCCTGGAGCTGTTCAAC | |

**Supplementary Material: Table 3: List of plasmids used for the study**

| Serial Number | Gene | Catalog name/Number | Company |
|---|---|---|---|
| 1 | USP1sh RNA | TRCN0000030769 | Sigma Aldrich Co. |
| 2 | pLKO scr | pLKO.1 puro<br>Plasmid:8453 | Addgene |
| 3 | USP1 over expression | Flag-HA-USP1<br>Plasmid: 22596 | Addgene |
| 4 | ID2sh RNA | TET-pLKO.1 PURO<br>shId2 #2<br>Plasmid: 83090 | Addgene |
| 5 | ID2 over expression | pLV-tetO-Id2<br>Plasmid: 70764 | Addgene |
